# Microglia drive demyelination via multiple sclerosis antibodies and BTK signaling

**DOI:** 10.64898/2026.08.25.747169

**Authors:** Lindsay A. Osso, Helena J. Barr, Michael E. Stockton, Maureen Wentling, Stephanie Karas, Rongchen Huang, Graham C. Peet, Katherine S. Given, Arlo Simmerman, Crystal R. McClain, Mohammad Mansoor, Dustin P. Fykstra, Kellie Horan, Clara Mutschler, Connon I. Thomas, Anza Darehshouri, Lan Lee, Ross C. Gruber, Dimitry Ofengeim, Anna Williams, Wendy B. Macklin, Gregory P. Owens, Jeffrey L. Bennett, Ethan G. Hughes

## Abstract

Microglia are the predominant immune cells in multiple sclerosis (MS) demyelinating lesions, where they phagocytose myelin, but whether they destroy myelin or merely scavenge its debris is unknown. Here, we explore whether pathogenic autoantibodies found in MS may induce the phagocytic destruction of myelin by microglia. Applying patient-derived, myelin-targeting antibodies to the mouse cortex, we developed an *in vivo* model of MS with focal demyelination that depended on epitope specificity and Fc gamma receptor and complement binding. Longitudinal monitoring of microglia-myelin interactions using *in vivo* two-photon microscopy revealed rapid microglial envelopment of intact myelin driving myelin loss, while single-cell RNA sequencing identified a demyelination-associated microglial signature. Parallel changes were observed in human MS lesions, where microglia enveloped intact myelin and similar genes were upregulated. Inhibition of Bruton’s tyrosine kinase (BTK) limited microglial transcriptional changes and prevented myelin loss following microglial envelopment. These findings directly implicate microglia in pathological myelin loss and support BTK inhibition as a therapeutic strategy to prevent demyelination by modulating microglia behavior.

---

Multiple sclerosis (MS) is a chronic autoimmune demyelinating disease of the central nervous system (CNS) that leads to neuronal injury and the accumulation of neurological impairment^1^. Current consensus supports demyelination in MS being induced by infiltrating T cells, with recent therapeutic successes in depleting non-antibody-secreting B cells also pointing to an important role for B cell cytokine secretion and antigen presentation^2^. However, the precise mechanisms leading to CNS demyelination remain unresolved.

Microglia, the resident macrophages of the CNS parenchyma, are the most prevalent immune cells in MS demyelinating lesions^3–5^. These cells are heavily enriched in areas of active inflammatory demyelination, including the core of active lesions and the borders of chronic active lesions^3–5^. It is well-established that microglia phagocytose myelin in these regions^4,6^, but whether microglia play an active role in myelin loss or merely uptake myelin debris is unclear.

Another hallmark feature of MS is the presence of cerebrospinal fluid (CSF) IgG antibodies^7^. These antibodies are produced by antigen-activated clonally expanded B cells^8^ in CSF, meninges, and brain parenchyma^9^ and can comprise autoantibodies that pathogenically target myelin^10,11^. Myelin-targeting antibodies are widespread across the MS population, with more than half of patients harboring autoantibodies against complexes containing the myelin protein PLP1^11^. Evidence suggests these autoantibodies may contribute to demyelination in MS by targeting myelin for phagocytic destruction by microglia. IgG antibodies and complement components are prevalent in actively demyelinating MS lesions where they colocalize with intact myelin sheaths as well as with myelin phagocytosed by microglia or other mononuclear phagocytes^3,12,13^. In mouse models, myelin-targeting autoantibodies derived from patient CSF B cells are pathogenic, inducing loss of myelin and oligodendrocytes in a complement-dependent manner^10,11^. Furthermore, targeting antibody production through depletion of antibody-secreting B cells substantially ameliorates the disease course in the MS model experimental autoimmune encephalomyelitis (EAE) as compared to depletion of non-antibody-secreting B cells alone^14^. However, the possibility that microglia mediate the demyelination caused by myelin autoantibodies has not yet been experimentally interrogated.

Microglial phagocytosis of antibody-targeted structures proceeds through two synergistic^15–17^ pathways: (1) microglial Fc gamma receptors binding to antibodies, and (2) microglial complement receptors binding to complement opsonins deposited on antibody-targeted cells^18,19^. Downstream of Fc gamma receptors, activation of Bruton’s tyrosine kinase (BTK)^20–22^ facilitates the phagocytosis of antibody-bound targets by mononuclear phagocytes^23–25^. The expression of this kinase is upregulated in mononuclear phagocytes in MS lesions^21,26^, though it has been pursued as a novel MS therapeutic target in large part for its role in activating B cells^27^. BTK inhibition decreases the appearance and expansion of lesions and limits the frequency of relapses in clinical trials^28–31^, and shows similar effects in EAE mouse models^22,26,32,33^. However, whether BTK inhibition might act to reduce demyelination by preventing microglial phagocytic destruction of myelin sheaths is unknown.

Real-time examination of microglial behavior throughout demyelination has been limited by the lack of focal and inducible immune-mediated models of MS that would facilitate *in vivo* monitoring of microglia-myelin interactions. Thus, building on previous work^10,11^, we developed a translationally relevant and treatment responsive *in vivo* model of immune-mediated demyelination by applying pathogenic MS patient-derived myelin-targeting antibodies and human complement-containing serum^10,11^ (MSrAb + hS) to the cortical surface of mice. Through a combination of longitudinal *in vivo* two-photon imaging and single-cell RNA sequencing, we characterized microglial dynamics throughout demyelination and their underlying transcriptional profiles. We found that microglia envelop intact myelin sheaths as early as fifteen minutes post-antibody application to induce demyelination, and upregulate a demyelinating genetic signature. Importantly, these changes are mirrored in MS patient lesions, where microglia envelop myelin sheaths and similar genes are upregulated. Testing the role of BTK inhibition in this process, we administered a brain-penetrant BTK inhibitor and found it limits microglial transcriptional changes and prevents the loss of myelin sheaths following their envelopment. These findings implicate microglia in myelin loss in MS and describe a novel mechanism by which BTK inhibition prevents demyelination.

## Results

### Cortical application of MS patient-derived recombinant antibodies induces rapid, focal oligodendrocyte loss

To assess the role of microglia in demyelination, we developed a new model of focal immune-mediated demyelination in the cortical gray matter. We used a pathogenic monoclonal recombinant antibody derived from MS patient CSF plasmablasts, which targets PLP1 complexes on CNS myelin^10,11,34^ (**Extended Data Fig. 1**). PLP1 complexes are targeted by intrathecally-produced autoantibodies in over half of MS patients tested^11^ and so understanding the mechanisms by which autoantibodies against these complexes lead to demyelination is of high clinical importance. Following surgical resection of the skull and dura over the motor cortex, we applied a cocktail containing patient-derived anti-PLP1 recombinant antibodies (MSrAb) and human complement-containing serum (hS) in sterile saline to the brain surface of mice expressing EGFP in mature, myelinating oligodendrocytes (*Mobp-EGFP*; **Fig. 1a**). We then used longitudinal *in vivo* two-photon imaging to track oligodendrocytes and their associated myelin sheaths under the site of application over several days (**Fig. 1a**). We have previously confirmed that this imaging approach faithfully reports changes to myelin sheath length and presence both via immunostaining and spectral confocal reflectance microscopy (SCoRe)^35^. Finally, we corroborated our *in vivo* findings through histological analysis of the region (**Fig. 1a**).

**Fig. 1:**
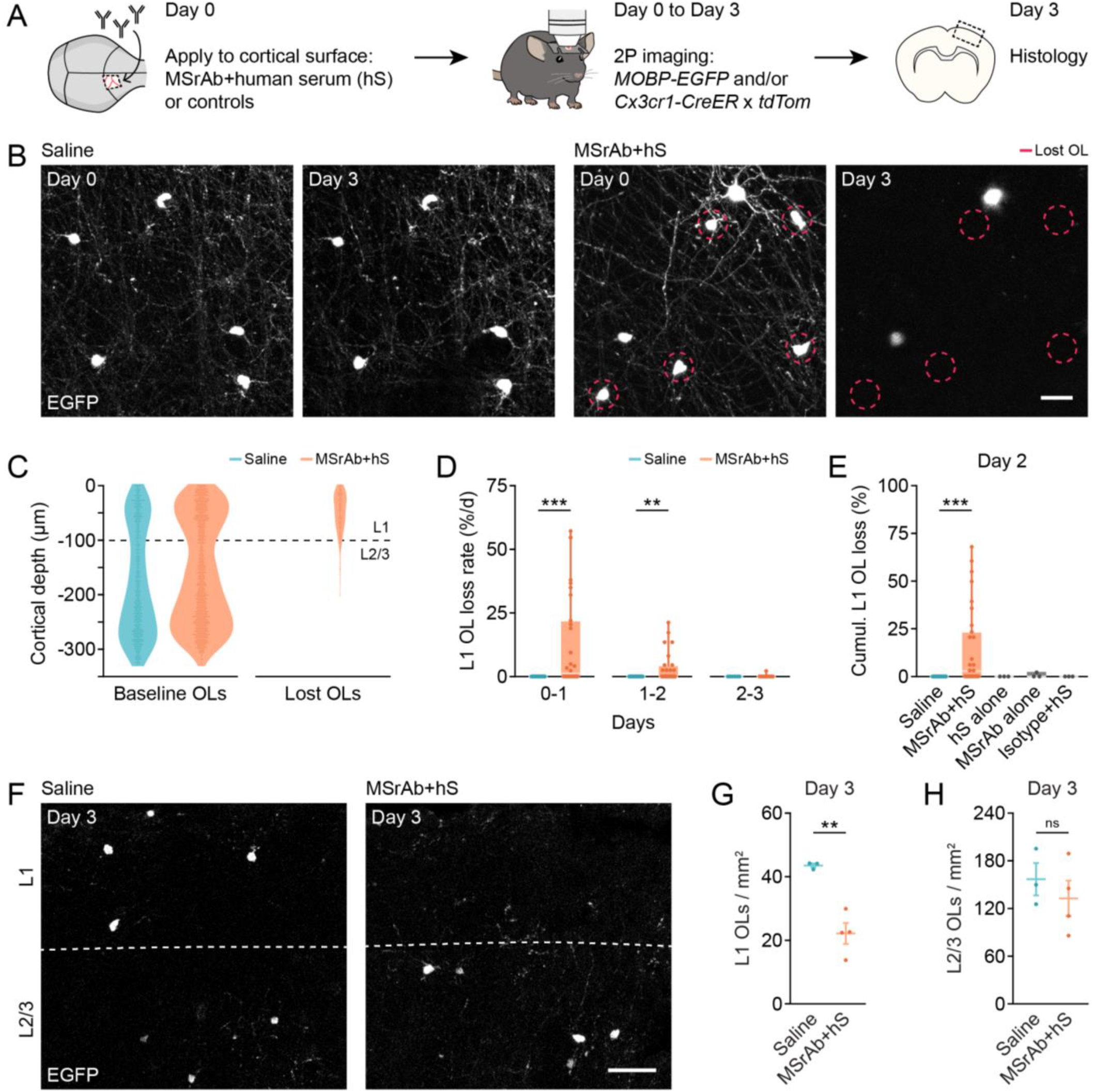
Cortical application of MS patient-derived recombinant antibodies induces rapid layer 1 oligodendrocyte loss in a complement-dependent manner. **(A)** Experimental timeline for antibody administration, two-photon *in vivo* imaging, and histology. **(B)** Representative images of EGFP+ oligodendrocytes on Day 0 and Day 3 from longitudinal *in vivo* images. Red circles indicate oligodendrocytes lost between Day 0 and Day 3 observed in MSrAb + hS treated mice. Scale bar is 20 μm. **(C)** Distribution by cortical depth of oligodendrocytes at baseline and those lost by Day 3 in Saline (n = 17) and MSrAb + hS (n = 28) treated mice indicates loss from MSrAb + hS treatment is biased to layer 1. **(D)** Rate of layer 1 oligodendrocyte loss (as a % of baseline oligodendrocytes) is elevated in MSrAb + hS treated mice on Days 0-1 and 1-2. **(E)** Cumulative loss of layer 1 oligodendrocytes (as a % of baseline oligodendrocytes) by Day 2 is elevated in MSrAb + hS treated mice, but not when MSrAb or hS are lacking or when an isotype antibody is used. **(F)** Representative images of histology for EGFP+ oligodendrocytes on Day 3 reveal loss of layer 1 oligodendrocytes in MSrAb + hS treated mice. Dashed line marks border of cortical layers, as determined by DAPI. Scale bar is 50 μm. **(G)** Reduction in layer 1 oligodendrocytes on Day 3 by histology in MSrAb + hS treated mice. **(H)** No loss of layer 2/3 oligodendrocytes on Day 3 by histology. In **D**, Kruskal-Wallis: Day 0-1 (ChiSquare (1) = 11.5274, p = 0.0007; Saline: n = 17, median (IQR) = 0 (0, 0); MSrAb + hS: n = 28, median (IQR) = 1.1364 (0, 21.5839)), Day 1-2 (ChiSquare (1) = 8.4227, p = 0.0037; Saline: n = 17, median (IQR) = 0 (0, 0); MSrAb + hS: n = 28, median (IQR) = 0 (0, 3.8929)), Day 2-3 (ChiSquare (1) = 0.5, p = 0.4795; Saline: n = 6, median (IQR) = 0 (0, 0); MSrAb + hS: 12, median (IQR) = 0 (0, 0). In **E**, Kruskal-Wallis (ChiSquare (4) = 19.3188, p = 0.0007; Saline: n = 18, median (IQR) = 0 (0, 0); MSrAb + hS: n = 30, median (IQR) = 2.8030 (0, 23.2202); hS alone: n = 3, median (IQR) = 0 (0, 0); MSrAb alone: n = 3, median (IQR) = 0 (0, 2.1505); Isotype + hS: n = 3, median (IQR) = 0 (0, 0). Post-hoc Dunn: MsrAb + hS vs. Saline (Z = 4.0168, p = 0.0006). In **G**, t test (t (5) = 5.4190, p = 0.0029; Saline: n = 3, mean ± SEM = 43.5483 ± 0.6429; MSrAb + hS: n = 4, mean ± SEM = 22.1830 ± 3.3010). In **H**, t test (t (5) = 0.7653, p = 0.4786; Saline: n = 3, mean ± SEM = 156.8847 ± 20.4375; MSrAb + hS: n = 4, mean ± SEM = 132.7518 ± 22.3910). ^*^ p < 0.05, ^**^ p < 0.01, ^***^ p < 0.001, n = mice; two-sided statistical tests. See **Extended Data Table 1** for statistical details. OL = oligodendrocyte.

Application of MSrAb + hS induced focal oligodendrocyte loss observed via longitudinal *in vivo* imaging (**Fig. 1b-e**) and histology (**Fig. 1f-h**). Loss was most prominent in cortical layer 1, closest to the site of MSrAb + hS application (**Fig. 1c, g-h**), reminiscent of subpial demyelinating lesions in MS patients^36^. Furthermore, oligodendrocyte loss was rapid: the vast majority (78%) occurred within the first day and was complete by the end of the second day (**Fig. 1d**).

Oligodendrocyte loss was never observed with saline application (**Fig. 1b-e**) nor human complement-containing serum application alone (**Fig. 1e**), and was negligible with MSrAb application alone (**Fig. 1e**), indicating loss was both antibody- and serum-dependent. Moreover, oligodendrocyte loss was not a non-specific response to antibody presence as an isotype control antibody (measles clone 2B4) + hS did not induce loss (**Fig. 1e**). Taken together, these findings indicate that MSrAb induces oligodendrocyte loss in an antibody-specific and serum-dependent manner.

### Cortical application of MS patient-derived recombinant antibodies induces targeted, rapid myelin sheath loss

In addition to loss of oligodendrocytes, we observed loss of myelin sheaths in MSrAb + hS-treated mice – both from oligodendrocytes that were lost (**Fig. 2a**) and from stable oligodendrocytes (**Fig. 2b**). We traced an unbiased subset of individual layer 1 sheaths from the day of surgery and tracked their morphological status and presence over time (**Fig. 2c**). MSrAb + hS-treated mice lost 54% (median, IQR 42-95) of their layer 1 sheaths by the second day, in stark contrast to saline-treated mice, which maintained stable sheaths (median 0% (IQR 0-0) sheath loss) (**Fig. 2d**). The magnitude of sheath loss was positively correlated with the magnitude of oligodendrocyte loss (p = 0.024, R^2^ = 0.54, Y = 0.99X + 45.2), with the highest levels of sheath loss occurring in mice that experienced oligodendrocyte death. Interestingly, all MSrAb + hS-treated mice experienced at least some sheath loss – even those that did not exhibit oligodendrocyte loss – and sheath loss was substantially higher than oligodendrocyte loss (**Fig. 2e**). Sheath loss independent of oligodendrocyte death was also observed histologically in MSrAb + hS-treated mice. Myelin levels were severely reduced in layer 1 (**Fig. 2f-g**) where oligodendrocytes were lost (**Fig. 1c, f-g**), but were also reduced in layer 2/3 (**Fig. 2f, h**) where oligodendrocytes were maintained (**Fig. 1c, f, h**). Sheath loss was not accompanied by axon degeneration via immunostaining (**Extended Data Fig. 2a-b**) and we observed that axons remain sufficiently healthy through the demyelinating insult to enable remyelination (**Extended Data Fig. 2c**), consistent with direct pathological targeting of sheaths. Like oligodendrocyte loss, myelin loss from MSrAb + hS application occurred rapidly, as observed via longitudinal *in vivo* imaging. 79% of total loss occurred over the first day, with diminishing rates of sheath loss thereafter (**Fig. 2i-j**). Thus, MSrAb + hS treatment led to substantial and rapid loss of myelin sheaths.

**Fig. 2:**
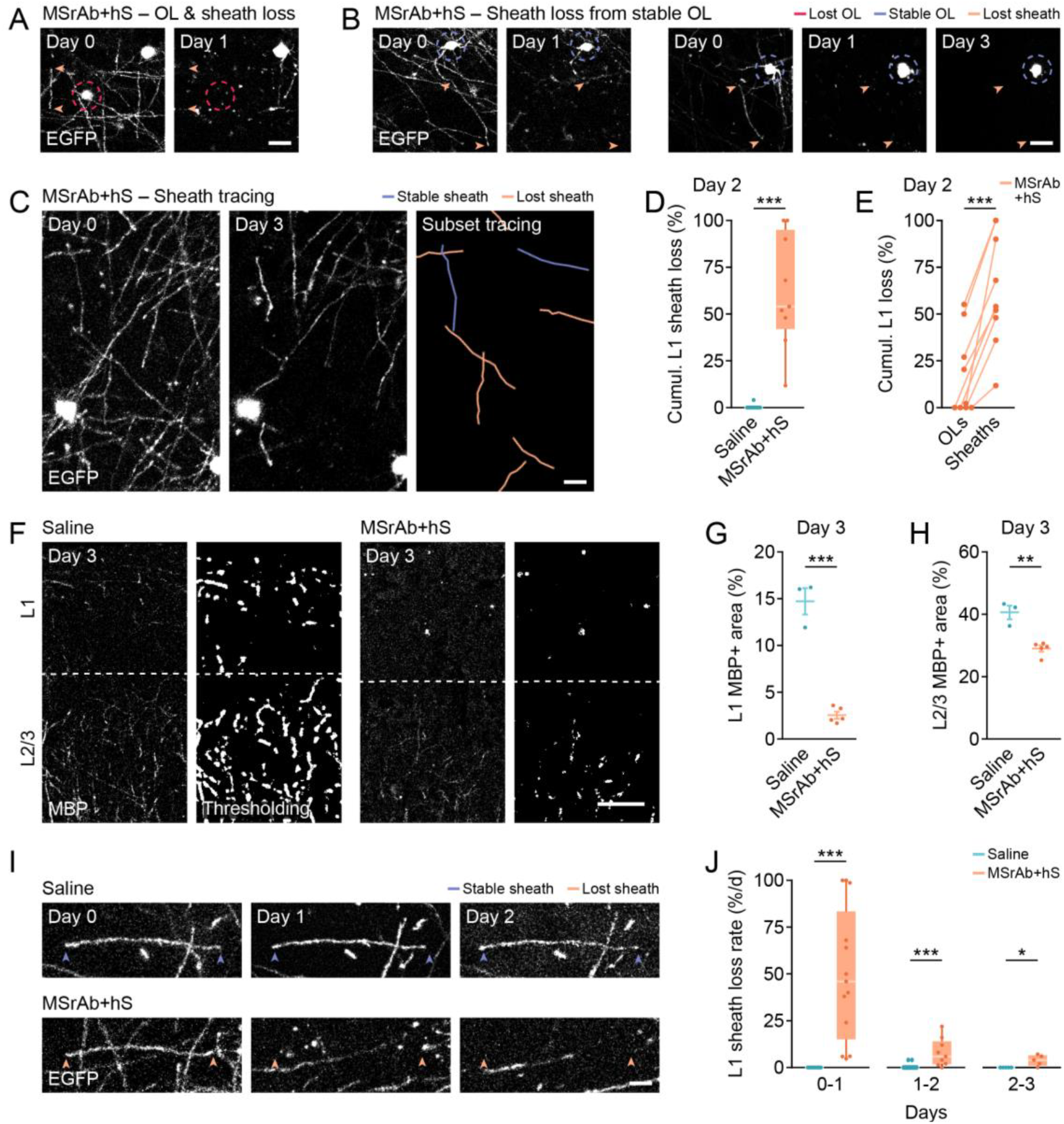
Rapid myelin loss induced by MS patient-derived recombinant antibodies. **(A-B)** Example longitudinal *in vivo* two-photon images of myelin loss with loss of the oligodendrocyte soma (A) and without loss of the oligodendrocyte soma (B). Orange arrowheads point to both termini of the myelin sheath. A red circle indicates a lost oligodendrocyte while blue circles indicate surviving, stable oligodendrocytes. Scale bar is 20 μm. **(C)** Example tracing of layer 1 myelin sheaths from longitudinal *in vivo* two-photon images showing stable sheaths (blue) and lost sheaths (orange). Unbiased tracing was performed on a subset of sheaths. Scale bar is 10 μm. **(D)** Cumulative loss of layer 1 sheaths (as a % of baseline sheaths) by Day 2 is elevated in MsrAb + hS treated mice. **(E)** MSrAb + hS treatment causes greater cumulative sheath loss than oligodendrocyte loss (both as a % of baseline levels) by Day 2. **(F)** Representative images of MBP immunostaining and thresholding analysis on Day 3 demonstrate loss of myelin in MSrAb + hS treated mice. Dashed line marks border of cortical layers, as determined by DAPI. Scale bar is 50 μm. **(G-H)** Reduction in layer 1 (G) and layer 2/3 (H) myelin area on Day 3 by histology in MSrAb + hS treated mice. **(I)** Example longitudinal *in vivo* two-photon images showing layer 1 sheath loss between Day 0 and 1 in MsrAb + hS treated mice. Arrowheads point to both termini of stable sheaths (blue) and lost sheaths (orange). Scale bar is 10 μm. **(J)** Rate of layer 1 sheath loss (as a % of baseline sheaths) is elevated in MSrAb + hS treated mice from Day 0 to Day 3. In **D**, Kruskal-Wallis (ChiSquare (1) = 18.2123, p < 0.0001; Saline: n = 13, median (IQR) = 0 (0, 0); MSrAb + hS: n = 9, median (IQR) = 54 (42, 95). In **E**, paired t test (t (8) = 6.6130, p = 0.0002, n = 9; OLs: mean ± SEM = 17.21 ± 7.485; Sheaths: mean ± SEM = 62.2 ± 10.05). In **G**, t test (t (6) = 10.7302, p < 0.0001; Saline: n = 3, mean ± SEM = 14.7221 ± 1.3966; MSrAb + hS: n = 5, mean ± SEM = 2.5542 ± 0.3718). In **H**, t test (t (6) = 5.6367, p = 0.0013; Saline: n = 3, mean ± SEM = 40.6014 ± 2.1936; MSrAb + hS: n = 5, mean ± SEM = 29.0037 ± 0.9684). In **J**, Kruskal-Wallis: Day 0-1 (ChiSquare (1) = 21.4551, p < 0.0001; Saline: n = 13, median (IQR) = 0 (0, 0); MSrAb + hS: n = 13, median (IQR) = 46 (15, 83.3421), Day 1-2 (ChiSquare (1) = 13.4863, p = 0.0002; Saline: n = 13, median (IQR) = 0 (0, 0); MSrAb + hS: n = 9, median (IQR) = 5.8824 (1.6579, 14), Day 2-3 (ChiSquare (1) = 5.5385, p = 0.0186; Saline: n = 5, median (IQR) = 0 (0, 0); MSrAb + hS: n = 5, median (IQR) = 3.9216 (1, 6.5211). ^*^ p < 0.05, ^**^ p < 0.01, ^***^ p < 0.001, n = mice; two-sided statistical tests. See **Extended Data Table 1** for statistical details. OL = oligodendrocyte.

### MS patient-derived recombinant antibodies alter microglial gene expression with similarities to MS patients

Microglia are the most prevalent immune cells in MS demyelinating lesions^3–5^, where they exhibit altered gene expression^37–40^. Thus, we next probed for changes in microglial gene expression in response to MS patient-derived recombinant antibodies via single-cell RNA sequencing. In mice expressing EGFP in cortical microglia (*Cx3cr1-EGFP*), we micro-dissected cortical tissue from beneath the area of MSrAb + hS application and from the intact contralateral cortex three days after surgery (**Fig. 3a**). Using fluorescence-activated cell sorting (FACS), we recovered live EGFP+ cells before capture using the 10X Chromium V3 platform (**Fig. 3a**). After quality control (**Extended Data Fig. 3a-c**), unsupervised clustering of 5855 total cells identified five populations (**Fig. 3b-c, Extended Data Table 2**). A “homeostatic” cluster comprised the majority of the total cell population **(Fig. 3c**). Also prevalent was a “demyelination” cluster that was heavily biased toward the MSrAb + hS-treated cortex (85.0% of cluster found in MSrAb + hS vs. 14.9% of cluster found in control) and made up a larger proportion of the microglia in MSrAb + hS cortex (31.3%) than in control cortex (5.1%) (**Fig. 3c**).

**Fig. 3:**
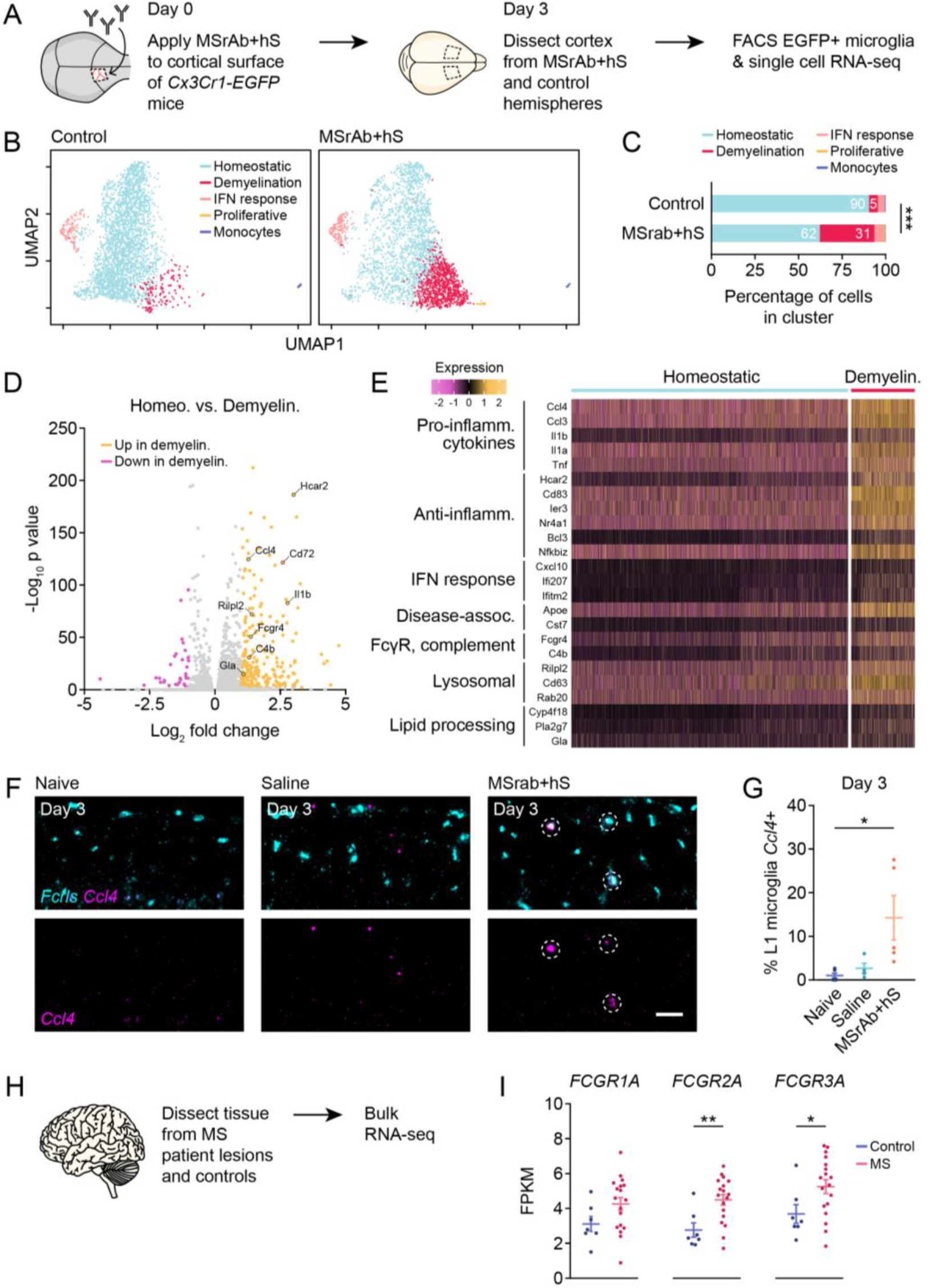
MS patient-derived recombinant antibodies induce transcriptional changes in microglia shared with MS patients. **(A)** Experimental timeline for antibody administration and scRNA-seq in n = 3 mice. **(B)** Unsupervised clustering distinguishes homeostatic, demyelination-associated, interferon-responsive, and proliferative microglia as well as monocytes in control and MSrAb + hS samples. **(C)** Proportion of microglia clusters differs between control and MSrAb + hS samples. **(D)** Volcano plot of distribution of upregulated (yellow) and downregulated (magenta) genes in demyelination vs. homeostatic microglia clusters. Colorized genes have an absolute fold change > 2 and adjusted p < 0.01. **(E)** Heat map of differently expressed genes between demyelination vs. homeostatic microglia clusters, selected from list of statistically significant genes. Expression range defined by Seurat normalized scaling method. **(F)** Representative images of RNAscope on Day 3 reveals upregulation of *Ccl4* in layer 1 microglia (*Fcrls*+) in MSrAb + hS treated mice. *Ccl4+Fcrls*+ microglia are outlined with a dashed circle. Scale bar is 50 μm. **(G)** Increase in % of layer 1 microglia expressing *Ccl4* on Day 3 by RNAscope in MSrAb + hS treated mice. **(H)** Experimental protocol for MS patient bulk RNA-seq. **(I)** Increase in high-affinity, activating Fc gamma receptor genes in MS patient tissue. In **C**, Chi-square (ChiSquare (4) = 749.2, p < 0.0001). In **G**, ANOVA (F (2, 11) = 5.1775, p = 0.0260; Naive: n = 5, mean ± SEM = 0.9146 ± 3.145; Saline: n = 4, mean ± SEM = 2.5796 ± 3.517; MSrAb + hS: n = 5, mean ± SEM = 14.2017 ± 3.145. Post-hoc Tukey’s: MsrAb + hS vs. Naive (p = 0.0306). In **I**, *FCGR1A*: t test (t (23) = 1.758, p = 0.0921; Control: n = 7, mean ± SEM = 3.114 ± 0.4234; MS: n = 18, mean ± SEM = 4.254 ± 0.3671). *FCGR2A*: t test (t (23) = 3.107, p = 0.0050; Control: n = 7, mean ± SEM = 2.761 ± 0.4072; MS: n = 18, mean ± SEM = 4.495 ± 0.3085). *FCGR3A*: t test (t (23) = 2.165, p = 0.0410; Control: n = 7, mean ± SEM = 3.688 ± 0.5334; MS: n = 18, mean ± SEM = 5.256 ± 0.3995). ^*^ p < 0.05, ^**^ p < 0.01, ^***^ p < 0.001 unless otherwise specified, n = mice; two-sided statistical tests. See **Extended Data Table 1** for statistical details. A full comparison of gene expression between clusters and between groups can be found in **Extended Data Tables 3 and 4**.

Compared to the homeostatic cluster, microglia in the demyelination cluster upregulated expression of numerous genes (**Fig. 3d**). A full comparison of gene expression between clusters and between groups can be found in **Extended Data Tables 3 and 4**, respectively. Specifically, demyelination-associated microglia were characterized by upregulated expression of genes involved in microglial inflammation, including pro-inflammatory cytokines (*Ccl4, Ccl3, Il1b, Il1a, Tnf*) and pro-inflammatory (*Cd72*) and anti-inflammatory (*Hcar2, Cd83, Ier3, Nr4a1, Bcl3, Nfkbiz*) modulators (**Fig. 3d-e**). Upregulation of interferon response gene expression was also apparent, both in the demyelina tion cluster (*Cxcl10, Ifi207, Ifitm2*) as well as in a third “interferon response” cluster (*Ifit3, Irf7, Usp18, Ifitm3*) (**Fig. 3b-c, e**). Several of these genes (*CCL4, IL1B, CD83, NR4A1, IFITM2)* are highly expressed in microglia in MS^38–41^, including the pro-inflammatory cytokine *CCL4*, which is upregulated by microglia in actively demyelinating MS lesions^42^. Thus, we sought to verify the upregulation of this cytokine in layer 1 microglia following MSrAb + hS treatment via RNAscope. Mirroring microglia in demyelinating MS lesions, the number of *Ccl4*-expressing layer 1 microglia was elevated in MSrAb + hS-treated mice but not in saline-treated or naïve mice (**Fig. 3f-g**). Additionally, we validated the increased expression of the inflammatory modulator *Cd72*^43^ in MsrAb + hS-treated layer 1 microglia (**Extended Data Fig. 3d-e**).

The demyelination-associated cluster shared several commonalities with previously described disease-associated microglial populations (DAMs)^44,45^, notably via common upregulation of genes including *Apoe, Cst7, Ccl4, and Ccl3* (**Fig. 3d-e**). In MS and particularly in actively demyelinating MS lesions, *APOE* is consistently upregulated in microglia across datasets^37–40^. Intriguingly, the demyelination-associated cluster showed upregulation of genes involved in antibody-dependent phagocytosis. We observed upregulation of the activating Fc gamma receptor *Fcgr4* as well as classical complement component and opsonin precursor *C4b*, whose cleavage product binds covalently to cellular targets to mark them for phagocytosis via recognition by phagocyte complement receptors (**Fig. 3d-e**). Consistent with myelin phagocytosis by microglia, this cluster was enriched for numerous genes involved in the lysosomal pathway (*Rilpl2, Cd63, Rab20*) and lipid processing (*Cyp4f18, Pla2g7, Gla*) (**Fig. 3d-e**). Analysis of bulk RNA sequencing datasets from post-mortem MS brain samples^26^ revealed similar upregulation of the activating Fc gamma receptors expressed by macrophages^46^ (**Fig. 3h-i**). *Fcgr4* does not have a direct analog in humans, but expression of the familial receptors *FCGR2A* and *FCGR3A* were significantly enhanced in MS patients relative to controls (**Fig. 3i**). Importantly, many of these genes (*FCGR2A, FCGR3A, CD63, RAB20, PLA2G7*) are upregulated in microglia in MS patients or in MS actively demyelinating lesions in other datasets^38–40^. Overall, these shared genetic signatures point to a common role for microglia in antibody recognition and subsequent cellular response within both MS patients and our animal model.

### Microglia envelop myelin following MS patient-derived recombinant antibody application reminiscent of MS patient lesions

Microglia are highly enriched at the borders of chronic active lesions and are thought to contribute to their slow expansion over time^5,38^. In these microglia-enriched actively demyelinating regions, intact myelin sheaths have been observed to be bound by IgG antibodies^12^, perhaps targeting them for demyelination via microglial phagocytosis. Thus, we sought to investigate the interactions between microglia and myelin sheaths in inflammatory demyelinating regions in MS. We collected post-mortem MS brain tissue containing chronic active lesions and stained for PLP1 (myelin) and IBA1 (mononuclear phagocytes, including microglia). In the microglia-enriched border of an occipital cortex white matter lesion (**Fig. 4a-b, Extended Data Fig. 4a**), we observed close interactions between IBA1+ cells and myelin, where IBA1+ cells appeared to envelop myelin sheaths (**Fig. 4c, Extended Data Fig. 4b, Extended Data Videos 1-3**). These intimate interactions were not observed in normal-appearing white matter of the same patient or in control brain tissue (**Extended Data Fig. 4c-d**).

**Fig. 4:**
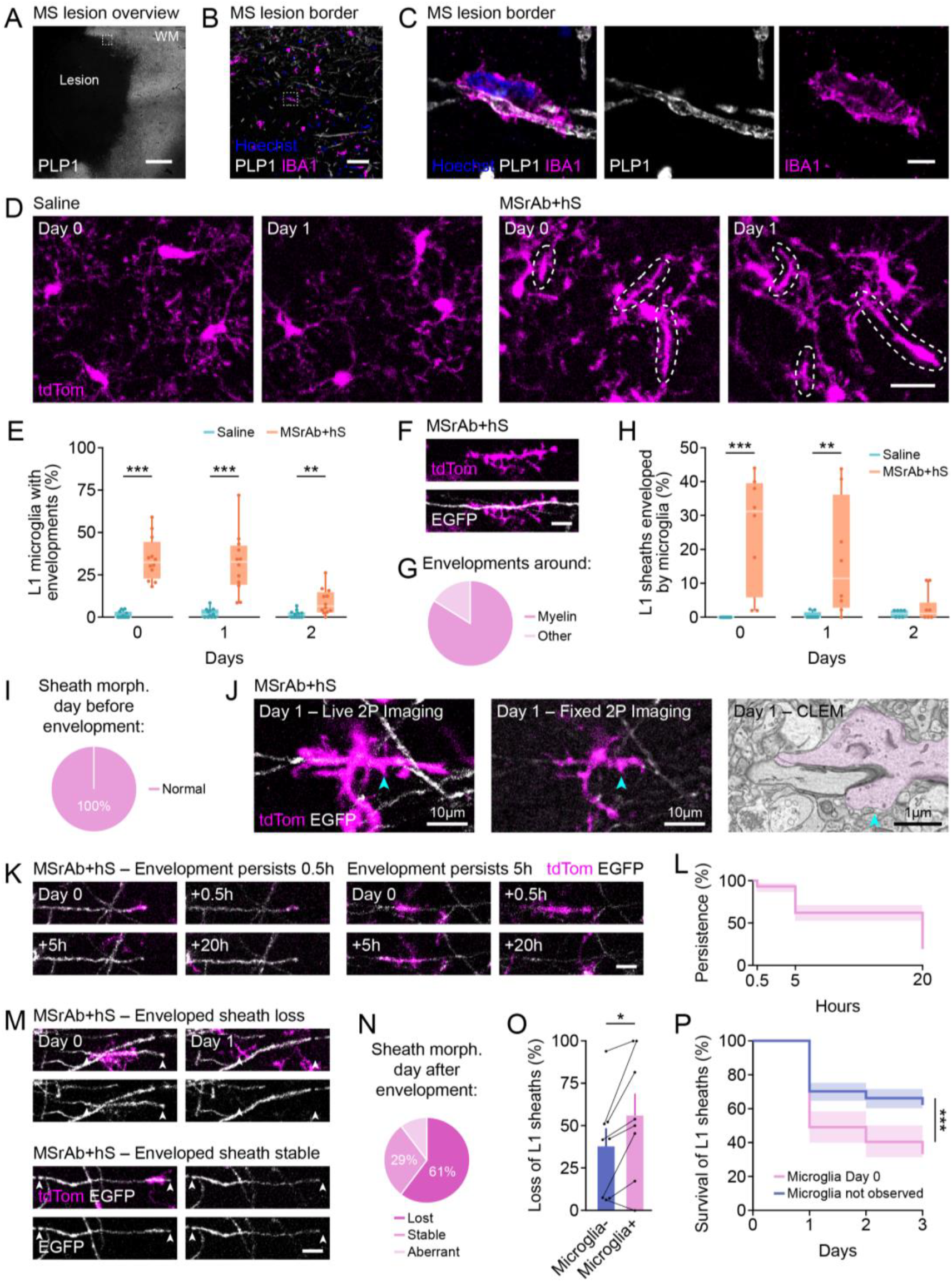
Rapid envelopment of sheaths by microglia in response to MS patient-derived recombinant antibodies leads to demyelination. **(A)** MS occipital cortical lesion area demarcated by PLP1 staining. A dashed square indicates the lesion border area in B. Scale bar is 1 mm. **(B)** IBA1+ cells (magenta) are enriched in the lesion border. A dashed rectangle outlines the IBA1+ cell in C. Scale bar is 50 μm. **(C)** Example 63X confocal image of an IBA1+ cell (magenta) enveloping a myelin sheath (white) in the lesion border. Scale bar is 5 μm. See **Extended Data Video 1** to view this microglia-myelin interaction through z. **(D)** Representative *in vivo* images of microglial envelopments (encircled) in MSrAb + hS treated mice on Days 0 and 1. Scale bar is 20 μm. **(E)** Increased % of layer 1 microglia forming envelopments in MSrAb + hS treated mice from Days 0-2. **(F)** Example *in vivo* image of a microglial envelopment (magenta) around a myelin sheath (white) in MSrAb + hS treated mouse. Scale bar is 10 μm. **(G)** The vast majority of microglial envelopments were around myelin (Day 0, n = 94 envelopments from 10 MSrAb + hS treated mice). **(H)** Increased % of layer 1 sheaths enveloped by microglia in MSrAb + hS treated mice on Days 0 and 1. **(I)** 100% of sheaths appeared normal the day prior to microglial envelopment (n = 25 sheaths from 7 MSrAb + hS mice). **(J)** *In vivo* image of a microglia envelopment of a myelin sheath on Day 1 before (left) and after (middle) fixation. Electron micrograph (right) of same microglia-sheath interaction shows close contact between enveloping microglial process and intact sheath. Arrowhead indicates same microglial structure between images. **(K)** Example longitudinal *in vivo* images of microglial envelopments (arrowheads) present immediately following surgery persisting for 0.5 h (top) and 5 h (bottom). Scale bar is 10 μm. **(L)** Persistence of microglia envelopments on sheaths over time following surgery (n = 103 sheaths from 3 MSrAb + hS mice). **(M)** Example longitudinal *in vivo* images of enveloped sheaths that were lost or remained stable. Arrowheads point to both termini of sheaths. Scale bar is 10 μm. **(N)** Proportion of sheaths lost (61%), stable (29%), or with aberrant morphology (10%) the day following microglial envelopment (n = 109 sheaths from 8 MSrAb + hS mice). **(O)** Sheaths enveloped by microglia were more likely to be lost than those never observed to be enveloped in our imaging timepoints. **(P)** Survival curve shows sheaths enveloped on Day 0 were more likely to be lost over time than those that were never observed to be enveloped in our imaging timepoints. In **E**, Day 0: Kruskal-Wallis (ChiSquare (1) = 16.7903, p < 0.0001; Saline: n = 11, median (IQR) = 0 (0, 3.2258); MSrAb + hS: n = 12, median (IQR) = 32.3793 (22.8656, 44.3078). Day 1: Kruskal-Wallis (ChiSquare (1) = 16.0833, p < 0.0001; Saline: n = 11, median (IQR) = 1.6949 (0, 4.5455); MSrAb + hS: n = 12, median (IQR) = 32.5598 (19.1483, 42.2947). Day 2: Kruskal-Wallis (ChiSquare (1) = 7.9798, p = 0.0047; Saline: n = 11, median (IQR) = 1.7391 (0, 2.8571); MSrAb + hS: n = 12, median (IQR) = 6.5080 (3.0386, 14.7166). In **H**, Day 0: Kruskal-Wallis (ChiSquare (1) = 12.9076, p = 0.0003; Saline: n = 8, median (IQR) = 0 (0, 0); MSrAb + hS: n = 8, median (IQR) = 31.1972 (5.9118, 39.5. Day 1: Kruskal-Wallis (ChiSquare (1) = 8.1250, p = 0.0044; Saline: n = 8, median (IQR) = 0 (0, 1.5); MSrAb + hS: n = 8, median (IQR) = 11.4583 (2.8054, 36.1111). Day 2: Kruskal-Wallis (ChiSquare (1) = 0.4730, p = 0.4916; Saline: n = 7, median (IQR) = 2 (0, 2); MSrAb + hS: n = 6, median (IQR) = 1.0417 (0, 4.2799). In **O**, paired t test (t (7) = 2.7780, p = 0.0274, n = 8 mice, Microglia negative: mean ± SEM = 37.75 ± 10.71, Microglia positive: mean ± SEM = 56 ± 12.89). In **P**, Log-Rank test (ChiSquare = 23.9552, p < 0.0001; Microglia Day 0: n = 69 sheaths, Microglia not observed: n = 98 sheaths). p < 0.05, ^**^ p < 0.01, ^***^ p < 0.001, n = mice unless otherwise specified; two-sided statistical tests. See **Extended Data Table 1** for statistical details. See **Extended Data Table 5** for human patient metadata.

We next sought to assess the cellular behavior of microglia in our mouse model during MSrAb-mediated demyelination. We bred triple transgenic mice (*Cx3cr1-CreER; Ai14; Mobp-EGFP*) to allow for the simultaneous visualization of myelinating oligodendrocytes (EGFP) and *Cx3cr1*-expressing myeloid lineage cells, including microglia (tdTomato). We then applied MSrAb + hS to the cortex and monitored microglia-myelin interactions *in vivo* in real-time using a combination of acute time-lapse and longitudinal imaging via two-photon microscopy (**Fig. 1a**). Cortical tdTomato-expressing microglial somas and characteristic ramified processes were readily visualized with this method (**Fig. 4d**, “Saline”, **Extended Data Fig. 5a**).

Microglia undergo changes in density and morphology upon their activation^47,48^. Therefore, we first investigated whether these phenotypes of cellular activation were impacted in our model. While we saw an increase in layer 1 microglial density and soma size following surgery, application of MSrAb + hS did not further alter these parameters (**Extended Data Fig. 5a-f**). However, we observed layer 1 microglia in MSrAb + hS-treated mice assume an unusual morphology (**Fig. 4d**). They rapidly extended their processes to form elongated protrusions, which were not observed in saline-treated mice **(Fig. 4d-e**). In examining our dual-color mice, we found that these protrusions were envelopments around myelin sheaths (**Fig. 4f-g**). These envelopments were generated incredibly rapidly in MSrAb + hS-treated mice, with 32.4% (median, IQR 22.9-44.3) of microglia possessing these structures on the day of surgery (mice imaged often less than 15 minutes following antibody application) (**Fig. 4e**), resulting in 31.2% (median, IQR 5.9-39.5) of sheaths being enveloped by microglia at this timepoint (**Fig. 4h**). Microglial envelopments were similarly prevalent the first day after surgery in MSrAb + hS-treated mice but tapered off by Day 2 (**Fig. 4d-e, h**). By contrast, microglia envelopments were not observed in a toxin-mediated model of cortical demyelination^49^ (**Extended Data Fig. 6a-d**).

Given the rapid onset of microglial envelopment, we suspected that sheaths were targeted by microglia prior to incurring damage. Indeed, sheaths enveloped by microglia appeared normal by *in vivo* imaging (**Fig. 4f**). Furthermore, when assessing sheaths enveloped by microglia beginning on Day 1 and beyond, we verified that 100% of these had a normal morphology at the prior *in vivo* imaging timepoint (**Fig. 4i, Extended Data Fig. 7a**). To obtain greater structural resolution of these microglia-myelin interactions, we performed correlative light and electron microscopy (CLEM) of envelopments identified in our *in vivo* images (**Fig. 4j**). Electron microscopy revealed an intimate interaction between microglia and myelin, with microglial processes directly contacting intact myelin sheaths along their length and circumference to form elaborate envelopments (**Fig. 4j**).

To investigate the stability of envelopments, we performed timelapse imaging immediately following surgery in MSrAb + hS-treated mice. These timelapses revealed that envelopments are dynamic structures, which could be observed growing or shrinking over time (**Extended Data Videos 4-5**). Tracking envelopments over 20 hours, we found that 93% of envelopments present at the beginning of imaging persisted over a half hour period (**Fig. 4k-l**). Five hours later, 62% of these envelopments remained, while by 20 hours, only 19% were still present (regardless of sheath status) (**Fig. 4k-l**). Meanwhile, we observed new envelopments form over this period, with 58% of envelopments present at 20 hours forming between five and 20 hours (n = 52 envelopments from n = 3 mice). Thus, microglial envelopments are dynamic structures that form immediately and in the hours following surgery and persist over several hours. Together, these results indicate that MS patient-derived antibodies and hS induce microglia to rapidly form and maintain envelopments over several hours around intact myelin, reminiscent of interactions in the actively demyelinating MS lesion border region.

### Microglia envelopments drive demyelination in response to MS patient-derived recombinant antibodies

Microglial envelopment of sheaths preceded sheath loss (**Fig. 2j** versus **Fig. 4h)**. Therefore, we explored whether microglia envelopment drives sheath loss during MSrAb-mediated demyelination. By tracking enveloped sheaths over time in MSrAb + hS-treated mice, we found that 61% of myelin sheaths enveloped by microglia were lost the day after envelopment (**Fig. 4m-n, Extended Data Video 5**). Meanwhile, 29% persisted as stable sheaths and the remainder assumed an aberrant morphology, including shrinking, blebbing, and recently-described myelin swelling^50^ (**Fig. 4m-n, Extended Data Fig. 7b-c**), suggesting that the presence of microglia envelopment is not itself sufficient to induce sheath loss. To test whether microglial envelopment drives sheath loss, we compared the fate of sheaths enveloped by microglia to sheaths that were not observed to be enveloped in our once-daily images within MSrAb + hS-treated mice. We found that sheaths enveloped at any timepoint were more likely to be lost than their non-enveloped counterparts from the same mouse (**Fig. 4o**). Furthermore, in tracking sheaths enveloped by microglia on the day of surgery, we observed they were less likely to survive than sheaths never observed to be enveloped (**Fig. 4p**).

These data align with the sequencing results and together suggest that sheaths are lost during MSrAb-mediated demyelination via microglial phagocytosis. Consistent with this hypothesis, we occasionally observed microglia envelopments persisting following the disappearance of the underlying myelin sheaths (**Extended Data Fig. 8a**). These persistent envelopments assumed a large, engorged morphology that covered where the sheath had been, suggesting they had recently phagocytosed the underlying sheath. Moreover, evidence of phagocytosis of *Mobp*-driven EGFP and myelin by microglia was apparent by *in vivo* imaging, electron microscopy, and histology (**Extended Data Fig. 8b-c**). CLEM revealed an EGFP inclusion in a microglia *in vivo* to be electron-dense myelin-like debris within the microglia by EM (**Extended Data Fig. 8b**). Meanwhile, immunostaining at the site of MSrAb + hS application showed microglia containing myelin within CD68+ lysosomes, indicative of phagocytosis (**Extended Data Fig. 8c**). Together, these results indicate that microglial envelopment of intact myelin drives demyelination in response to MS patient-derived recombinant antibodies likely via phagocytosis.

### MS patient-derived antibodies require an intact Fc region for Fc gamma receptor and complement binding to induce microglia-mediated demyelination

Microglia phagocytose antibody-targeted structures through two synergistic pathways: (1) microglial Fc gamma receptors binding to antibodies, and (2) microglial complement receptors binding to complement opsonins deposited on antibody-targeted cells following Fc-binding of C1q^18,19^. Our initial experiments indicate that complement-containing human serum is important to induce pathology in this model (**Fig. 1e**). To further examine these processes in microglia-mediated demyelination, we introduced two point mutations into the Fc region of the MSrAb (Mutant Fc), L234A and L235A (**Fig. 5a**). These mutations, commonly referred to as LALA, maintain antibody binding to antigen^51^ but block both subsequent Fc gamma receptor and complement C1q binding to the antibody Fc region^51,52^ (**Fig. 5a**) to prevent downstream responses including antibody-dependent phagocytosis^52^. To assess how this Fc mutation impacted microglial envelopment and demyelination in our model, we applied either MSrAb + hS or Mutant Fc + hS to the cortical surface of *Cx3cr1-CreER; Ai14; MOBP-EGFP* mice and performed longitudinal *in vivo* two-photon imaging of microglia and myelin.

**Fig. 5:**
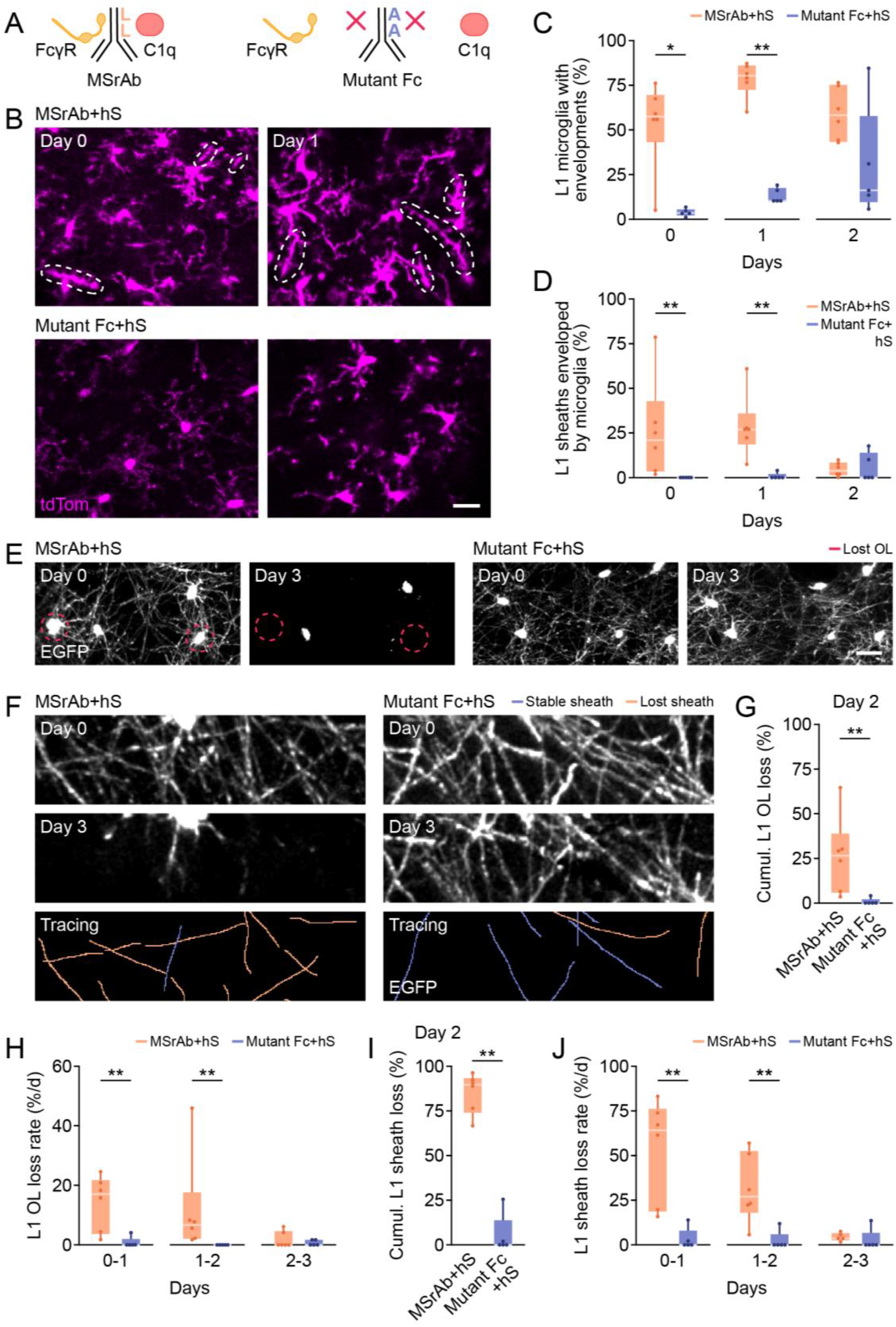
Intact Fc region required for MS patient-derived recombinant antibodies to induce microglial envelopments and demyelination. **(A)** LALA mutation in antibody Fc region prevents Fc gamma receptor and C1q binding. **(B)** Representative *in vivo* images of microglia envelopments (encircled) in MSrAb + hS-but not Mutant Fc + hS-treated mice on Days 0 and 1. Scale bar is 20 μm. **(C)** Rapid formation of envelopments in layer 1 microglia was prevented in Mutant Fc+ hS treated mice. **(D)** Rapid microglial envelopment of myelin sheaths was prevented in Mutant Fc + hS-treated mice. **(E)** Representative longitudinal *in vivo* images of EGFP+ oligodendrocytes on Day 0 and Day 3. Red circles indicate oligodendrocytes lost between Day 0 and Day 3 observed in MSrAb + hS treated mice. Scale bar is 25 μm. **(F)** Example longitudinal *in vivo* images of EGFP+ myelin on Day 0 and Day 3 with tracing of layer 1 myelin sheaths showing stable sheaths (blue) and lost sheaths (orange). Scale bar is 10 μm. **(G)** Cumulative loss of layer 1 oligodendrocytes (as a % of baseline oligodendrocytes) by Day 2 is prevented in Mutant Fc + hS-treated mice. **(H)** Reduced rate of layer 1 oligodendrocyte loss (as a % of baseline oligodendrocytes) in Mutant Fc + hS-treated mice relative to MsrAb + hS treated mice between Days 0-1 and 1-2. **(I)** Cumulative loss of layer 1 sheaths (as a % of baseline sheaths) by Day 2 is prevented in Mutant Fc + hS treated mice. **(J)** Reduced rate of layer 1 sheath loss (as a % of baseline sheaths) in Mutant Fc + hS treated mice relative to to MsrAb + hS treated mice on Days 0-1 and 1-2. In **C**, Kruskal-Wallis: Day 0 (ChiSquare (1) = 6.5333, p = 0.0106; MSrAb + hS: n = 6, median (IQR) = 52.2746 (42.7656, 69.4912); Mutant Fc + hS: n = 5, median (IQR) = 3.3058 (1.9854, 5.4701)), Day 1 (ChiSquare (1) = 7.5000, p = 0.0062; MSrAb + hS: n = 6, median (IQR) = 80.2365 (72.4324, 85.9304); Mutant Fc + hS: n = 5, median (IQR) = 10.6667 (9.9608,17.5075)), Day 2 (ChiSquare (1) = 2.7000, p = 0.1003; MSrAb + hS: n = 6, median (IQR) = 58.20793 (43.2615, 75.1412); Mutant Fc + hS: n = 5, median (IQR) = 16.0305 (9.5100, 57.6768)). In **D**, Kruskal-Wallis: Day 0 (ChiSquare (1) = 8.2500, p = 0.0041; MSrAb + hS: n = 6, median (IQR) = 20.8333 (3.4041, 20.8333); Mutant Fc + hS: n = 5, median (IQR) = 0 (0,0)), Day 1 (ChiSquare (1) = 7.8947, p = 0.0050; MSrAb + hS: n = 6, median (IQR) = 26.9231 (18.5185, 35.7843); Mutant Fc + hS: n = 5, median (IQR) = 0 (0,0)), Day 2 (ChiSquare (1) = 0.0786, p = 0.7792; MSrAb + hS: n = 6, median (IQR) = 3.8462 (1.3889, 8.3333); Mutant Fc + hS: n = 5, median (IQR) = 0 (0, 13.8235)). In **G**, Kruskal-Wallis: (ChiSquare (1) = 6.8444, p = 0.0089; MSrAb + hS: n = 6, median (IQR) = 26.4254 (5.7685, 38.8001); Mutant Fc + hS: n = 5, median (IQR) = 0 (0, 2.0408)). In **H**, Kruskal-Wallis: Day 0-1 (ChiSquare (1) = 6.8444, p = 0.0089; MSrAb + hS: n = 6, median (IQR) = 17.0411 (3.6995, 21.7571); Mutant Fc + hS: n = 5, median (IQR) = 0 (0, 2.0408)), Day 1-2 (ChiSquare (1) = 8.2500, p = 0.0041; MSrAb + hS: n = 6, median (IQR) = 6.7776 (2.0690,17.8354); Mutant Fc + hS: n = 5, median (IQR) = 0 (0,0)), Day 2-3 (ChiSquare (1) = 0.0447, p = 0.8325; MSrAb + hS: n = 6, median (IQR) = 0 (0, 4.6494); Mutant Fc + hS: n = 5, median (IQR) = 0 (0,1.6903)). In **I**, Kruskal-Wallis: (ChiSquare (1) = 7.6389, p = 0.0057; MSrAb + hS: n = 6, median (IQR) = 89.6368 (74.0196, 93.3048); Mutant Fc + hS: n = 5, median (IQR) = 0 (0, 13.7255)). In **J**, Kruskal-Wallis: Day 0-1 (ChiSquare (1) = 7.6389, p = 0.0057; MSrAb + hS: n = 6, median (IQR) = 64.4231 (18.6275, 76.3889); Mutant Fc + hS: n = 5, median (IQR) = 0 (0, 7.8431)), Day 1-2 (ChiSquare (1) = 6.8440, p = 0.0089; MSrAb + hS: n = 6, median (IQR) = 26.9231 (18.0556, 52.4510); Mutant Fc + hS: n = 5, median (IQR) = 0 (0, 5.8824)), Day 2-3 (ChiSquare (1) = 2.6129, p = 0.1060; MSrAb + hS: n = 6, median (IQR) = 3.8462 (2.7778, 6.7873); Mutant Fc + hS: n = 5, median (IQR) = 0 (0, 6.8627)). ^*^ p < 0.05, ^**^ p < 0.01, ^***^ p < 0.001, n = mice; two-sided statistical tests. See **Extended Data Table 1** for statistical details. OL = oligodendrocyte.

In contrast to MSrAb + hS-treated mice, microglial envelopments of myelin sheaths were almost entirely eliminated on Day 0 and Day 1 following surgery in Mutant Fc + hS-treated mice (**Fig. 5b-d**). By Day 2, the groups were indistinguishable, with envelopments forming in a small subset of microglia (median 16.0%, IQR 9.5-57.7) and around a negligible number of sheaths (median 0%, IQR 0-13.8) in Mutant Fc + hS-treated mice (**Fig. 5c-d**). Thus, intact binding of Fc receptors and/or C1q to the antibody Fc region is required for the rapid microglial envelopment of sheaths in response to MS patient-derived recombinant antibodies.

Given the importance of microglial envelopment to sheath loss (**Fig. 4o-p**) and the impaired neutralization of targets with this Fc mutation^51,52^, we hypothesized that oligodendrocytes and myelin would be spared in Mutant Fc + hS-treated mice. Indeed, we found that MS patient-derived recombinant antibodies were unable to induce oligodendrocyte or myelin loss when they lacked an intact Fc region (**Fig. 5e-j**). Layer 1 oligodendrocyte loss was essentially eliminated in Mutant Fc + hS-treated mice (**Fig. 5e, g-h**). Myelin sheaths were also preserved, with a median cumulative sheath loss of 0% (IQR 0-13.7) by Day 2 post-application in Mutant Fc + hS-treated mice as compared to the near complete sheath loss in MSrAb + hS-treated mice (median 96.1%, IQR 85.5-99.0) at the same timepoint (**Fig. 5f, i-j**). Together, these data demonstrate the importance of Fc-initiated signaling in inducing microglial envelopment and subsequent loss of myelin sheaths in response to MS patient-derived recombinant antibodies.

### Inhibition of BTK signaling attenuates microglial transcriptional changes from MS patient-derived recombinant antibodies

Given the striking preservation of oligodendrocytes and myelin in the absence of Fc gamma receptor and complement signaling, we next sought to investigate the molecular mechanisms downstream of these pathways driving microglia-mediated demyelination. Bruton’s tyrosine kinase (BTK) activation in microglia occurs downstream of Fc gamma receptor signaling^20–22^, and small interfering RNA knockdown or genetic deletion of this kinase prevents Fc gamma receptor-dependent phagocytosis of antibody-bound targets by mononuclear phagocytes^23–25^. In EAE, BTK signaling is partly responsible for the transcriptional changes that occur in microglia and contributes to worsening disease course^22,26^. BTK inhibition has emerged as a novel therapeutic strategy in MS, with recent clinical trials reporting positive effects in relapsing and some progressive patients^28–31,53^. However, how BTK inhibition impacts microglia-driven pathology in MS is incompletely understood. Thus, we sought to test whether we could prevent microglial transcriptional activation and demyelination in response to MS patient-derived antibodies via pharmacological inhibition of BTK. Notably, within the brain, expression of this kinase is restricted to microglia^54^.

We administered a brain-penetrant BTK inhibitor (BTKi), PRN2675, an analog of tolebrutinib, via oral gavage to mice expressing EGFP in microglia (*Cx3cr1-EGFP*; **Fig. 6a**). Treatment was administered daily for three days starting one day prior to MsrAb + hS application. To test the effects of BTK inhibition on microglial gene expression, we micro-dissected cortical tissue from beneath the area of MSrAb application and collected microglia for single-cell RNA sequencing (**Fig. 6a**). We compared gene expression in microglia from MSrAb + hS-treated hemispheres with and without BTK inhibition. Unsupervised clustering revealed homeostatic, demyelination-associated, and IFN responsive microglia as well as monocytes (**Fig. 6b, Extended Data Table 2**). Importantly, BTKi-treated mice had a substantially reduced proportion of demyelination-associated microglia (**Fig. 6c**). Comparing unclustered microglia, BTK inhibition led to the downregulation of numerous genes (**Fig. 6d**). In particular, we found downregulation of pro-inflammatory (*Ccl4, Ccl3, Il1b, Il1a, Tnf*), anti-inflammatory (*Hcar2, Cd83, Ier3, Nr4a1, Bcl3, Nfkbiz*), lysosomal (*Rilpl2*), and lipid processing (*Gla*) genes (**Fig. 6d-e**) that were upregulated with MSrAb + hS treatment (**Fig. 3d-e**). For a full comparison of gene expression between groups, see **Extended Data Table 6**. Interestingly, we did not observe changes in expression of *Fcgr4* or *C4b* with BTK inhibition (**Fig. 6d-e**), suggesting the early cellular responses to antibody are not impacted. Taken together, these data show that inhibiting BTK signaling changes the transcriptional profile of microglia responding to MS patient-derived antibodies by preventing the upregulation of inflammatory and phagocytic pathways.

**Fig. 6:**
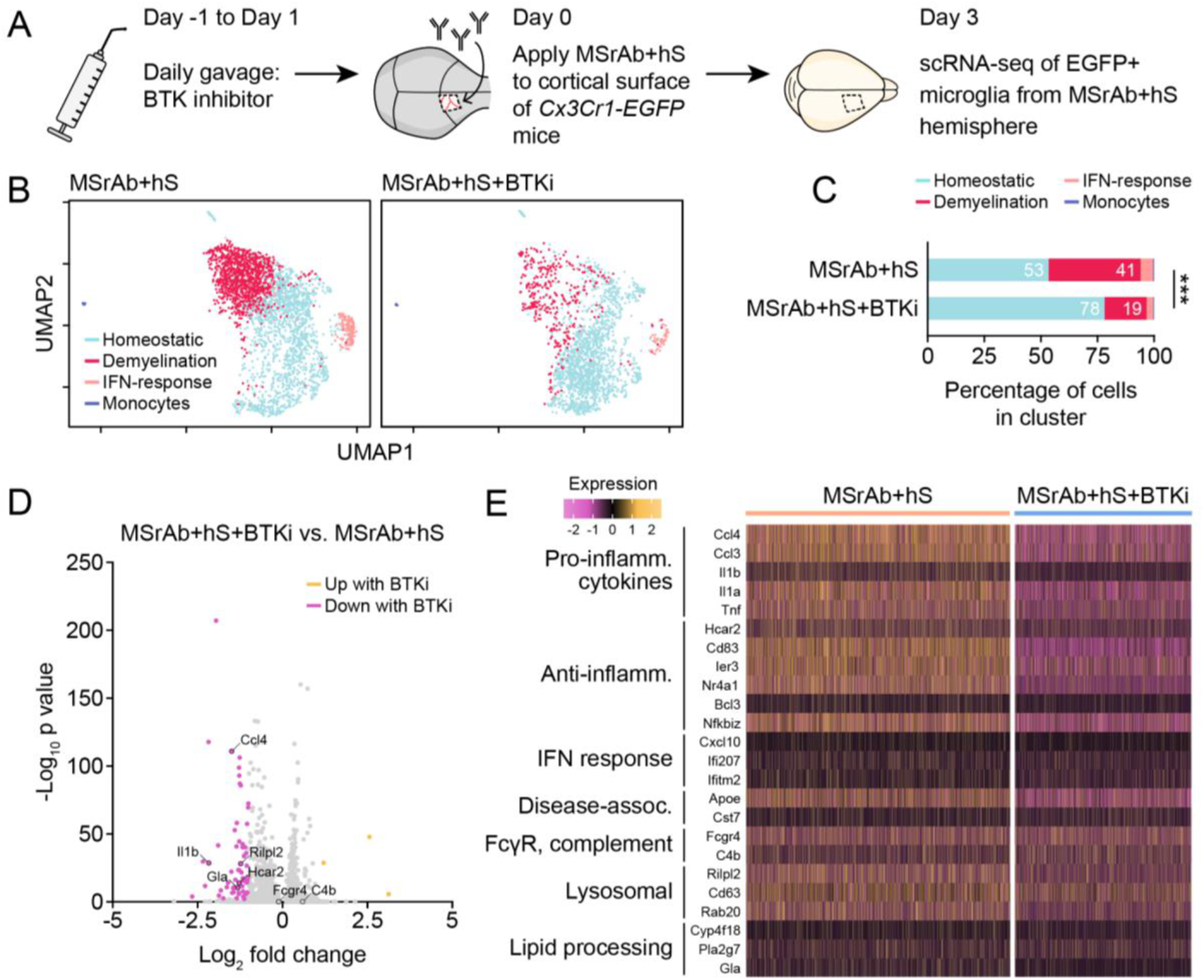
BTK inhibition limits upregulation of demyelination-associated genes in microglia. **(A)** Experimental timeline for BTK inhibitor and antibody administration and scRNA-seq in n = 3 mice. **(B)** Unsupervised clustering distinguishes homeostatic, demyelination-associated, and interferon-responsive microglia as well as monocytes in MSrAb + hS and MSrAb + hS + BTKi samples. **(C)** Proportion of microglia clusters differs between MSrAb + hS and MSrAb + hS + BTKi samples. Chi-square (ChiSquare (3) = 298.9, p < 0.0001). **(D)** Volcano plot of distribution of upregulated (yellow) and downregulated (magenta) genes in MSrAb + hS vs. MSrAb + hS + BTKi microglia samples. Colorized genes have an absolute fold change > 2 and adjusted p < 0.01. **(E)** Heat map of differential gene expression between MSrAb + hS + BTKi vs. MSrAb + hS microglia samples. Same genes from Fig. 3E. Expression range defined by Seurat normalized scaling method. ^*^ p < 0.05, ^**^ p < 0.01, ^***^ p < 0.001 unless otherwise specified, n = mice; two-sided statistical tests. See **Extended Data Table 1** for statistical details. A full comparison of gene expression between groups can be found in **Extended Data Table 6**.

### BTK inhibition reduces sheath loss following microglial envelopment to prevent antibody-mediated demyelination

Given the attenuation of MSrAb-mediated microglial gene expression changes with BTK inhibition, we next assessed the impact of inhibiting BTK on microglial morphology and cellular behavior. We compared *Cx3cr1-CreER; Ai14; Mobp-EGFP* mice treated with BTK inhibitor or vehicle and observed microglia and myelin via longitudinal *in vivo* imaging following the application of MSrAb + hS (**Fig. 7a**).

Similar to other conditions, layer 1 microglia density and soma size were increased, and there were no changes in morphological complexity by two days post-surgery, with no impact of BTKi on this phenomenon (**Extended Data Fig. 5g-i**). Next, we examined the occurrence of sheath envelopment by microglia post-MSrAb + hS application. BTKi-treated mice were equally likely to have microglia forming envelopment structures (**Fig. 7b-c**) as well as myelin sheaths enveloped by microglia (**Fig. 7d**) as vehicle-treated mice, both on the day of surgery and the following day. However, the outcome of enveloped sheaths differed radically between groups. While 71% of enveloped sheaths were lost the day after envelopment in MSrAb + hS + vehicle-treated mice, BTKi treatment reduced this proportion to 31% (**Fig. 7e**), consistent with the documented role of BTK in mediating Fc gamma receptor-mediated phagocytosis of antibody-bound targets^23–25^.

**Fig. 7:**
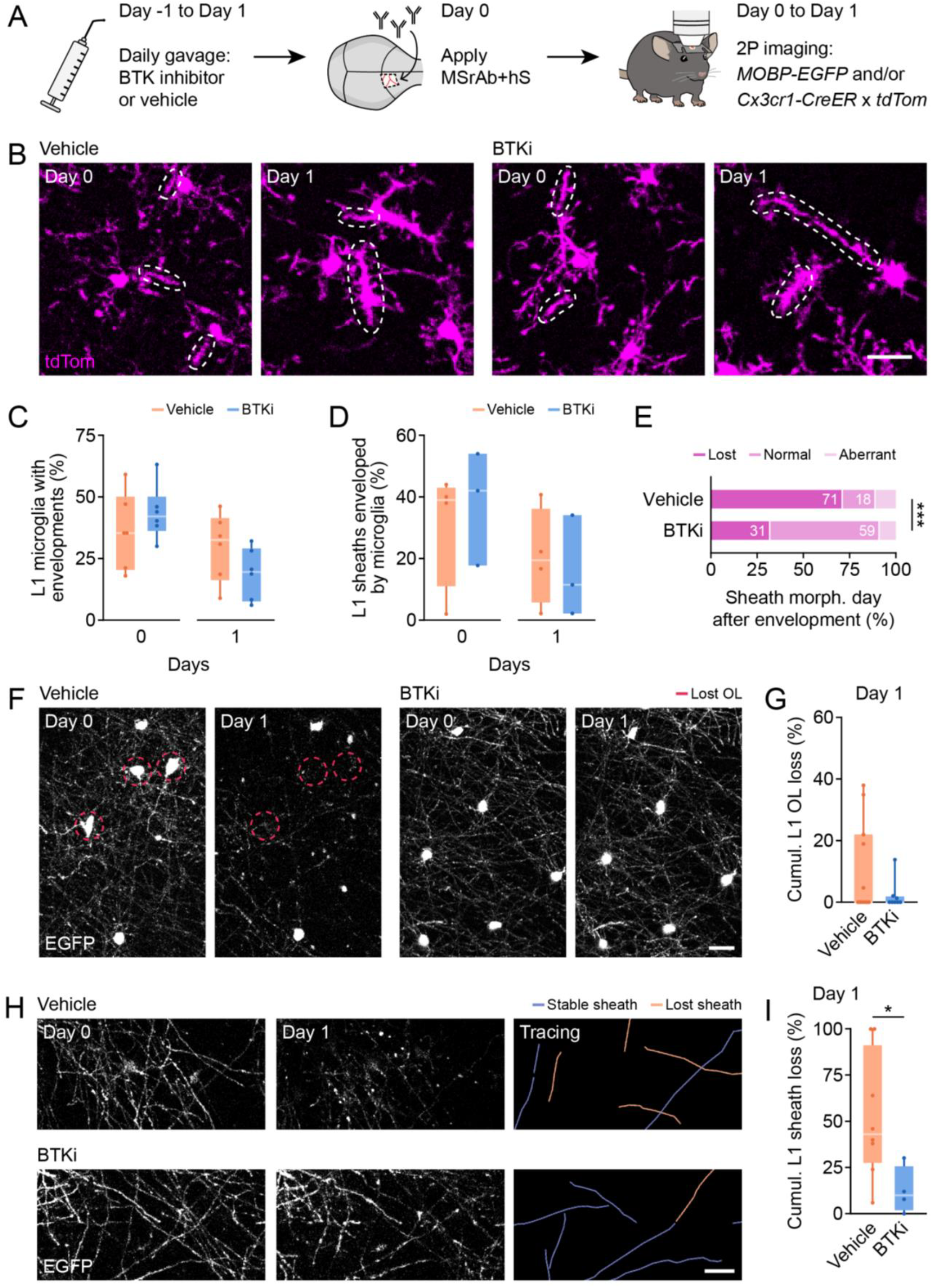
BTK inhibition prevents sheath loss following microglial envelopment. **(A)** Experimental timeline for BTK inhibitor and antibody administration and two-photon *in vivo* imaging. **(B)** Representative *in vivo* images of layer 1 microglia envelopments (encircled) in MSrAb + hS mice treated with Vehicle or BTKi on Days 0 and 1. Scale bar is 20 μm. **(C-D)** BTK inhibition does not affect % of layer 1 microglia forming envelopments (C) or % of layer 1 sheaths enveloped by microglia (D) on Days 0 or 1. **(E)** Proportion of sheaths lost following microglia envelopment is decreased by BTK inhibition. **(F)** Representative longitudinal *in vivo* images of EGFP+ oligodendrocytes on Day 0 and Day 1. Red circles indicate oligodendrocytes lost between Day 0 and Day 1. Scale bar is 20 μm. **(G)** Lack of high levels of layer 1 oligodendrocyte loss (as a % of baseline layer 1 oligodendrocytes) by Day 1 with BTK inhibition. **(H)** Example tracing of layer 1 myelin sheaths from longitudinal *in vivo* two-photon images showing reduced lost sheaths (orange) and increased stable sheaths (blue) with BTK inhibition. Scale bar is 20 μm. **(I)** Loss of layer 1 sheaths (as a % of baseline layer 1 sheaths) by Day 1 is prevented by BTK inhibition. In **C**, Kruskal-Wallis: Day 0 (ChiSquare (1) = 0.9231, p = 0.3367; Vehicle: n = 6, median (IQR) = 35.4012 (20.4173, 50.1488); BTKi: n = 6, median (IQR) = 41.9921 (36.1144,50.1645)); Day 1 (ChiSquare (1) = 2.3222, p = 0.1275; Vehicle: n = 6, median (IQR) = 32.5598 (16.176, 41.3244); BTKi: n = 6, median (IQR) = 19.5630 (7.6076, 29.0198)). In **D**, Kruskal-Wallis: Day 0 (ChiSquare (1) = 0.5000, p = 0.4795; Vehicle: n = 4, median (IQR) = 39 (11, 43); BTKi: n = 3, median (IQR) = 42 (17.6471, 54)); Day 1 (ChiSquare (1) = 0.2864, p = 0.5926; Vehicle: n = 4, median (IQR) = 19.4444 (5.7624, 36.1111); BTKi: n = 3, median (IQR) = 11.4286 (2.1277, 34.0909)). In **E**, Chi-square (ChiSquare (2) = 19.93, p < 0.0001). Vehicle: n = 62 sheaths from 4 mice; BTKi: n = 44 sheaths from 3 mice. In **G**, Kruskal-Wallis (ChiSquare (1) = 0.8449, p = 0.3580; Vehicle: n = 11, median (IQR) = 0 (0, 21.9512); BTKi: n = 8, median (IQR) = 0 (0, 1.8596)). In **I**, Kruskal-Wallis (ChiSquare (1) = 4.1684, p = 0.0412; Vehicle: n = 8, median (IQR) = 43 (27.5, 91); BTKi: n = 4, median (IQR) = 9.9216 (1.9608, 25.5)). p < 0.05, ^**^ p < 0.01, ^***^ p < 0.001, n = mice unless otherwise specified; two-sided statistical tests. See **Extended Data Table 1** for statistical details.

Lastly, we sought to examine whether BTK inhibition led to a reduction in oligodendrocyte and myelin sheath loss following MSrAb + hS administration. Mice treated with BTKi had a trend towards attenuated oligodendrocyte loss (**Fig. 7f-g**) and had substantially reduced myelin sheath loss, with BTKi-treated mice losing only 9.9% (median, IQR 2.0-25.5) of sheaths by Day 1 compared to 43.0% (median, IQR 27.5-91.0) of sheaths lost in vehicle-treated mice (**Fig. 7h-i**). Taken together, these results show that oral administration of a brain-penetrant BTK inhibitor reduces myelin loss following microglial envelopment to protect myelin in response to MS patient-derived myelin-targeting antibodies.

## Discussion

In this study, we find that microglia contribute to demyelination in response to MS autoantibodies through rapid myelin sheath envelopment and subsequent loss. This work adds to the emerging framework of how microglia and other brain-resident or infiltrating macrophages cause degeneration in MS, which may also involve the release of pro-inflammatory cytokines and reactive oxygen species^55,56^ as well as the phagocytosis of synapses^57^. Specifically, our data point to a role for microglia in causing phagocytic destruction of myelin through the engagement of the Fc gamma receptor, complement, and BTK signaling pathways (**Extended Data Fig. 9a**). First, microglia recognize and rapidly envelop antibody-bound myelin, dependent on the antibody Fc region. When the Fc region of MSrAb was mutated to prevent Fc gamma receptor and complement component C1q binding, we observed a profound suppression of sheath envelopment by microglia (**Fig. 5b-d**). Second, some enveloped sheaths are then removed. Envelopment increased the likelihood of sheath loss (**Fig. 4o-p**) and intervention to inhibit microglial envelopment also prevented demyelination (**Fig. 5e-j**), indicating the importance of envelopment to demyelination. However, envelopment alone was not sufficient for loss, as only just over half of enveloped sheaths were lost (**Fig. 4o**). Indeed, the transition from sheath envelopment to loss depends on BTK signaling, as inhibiting BTK reduced loss of enveloped sheaths by 2.5-fold (**Fig. 7e**), consistent with previous work demonstrating a role for BTK in facilitating Fc gamma receptor-mediated phagocytosis of antibody-bound targets^23–25^. As further evidence supporting microglial phagocytosis, we observed microglial phagocytic-like structures persisting after the disappearance of the underlying sheath, myelin debris in microglia, and *Mobp*-driven EGFP in microglial lysosomes (**Extended Data Fig. 8**).

While microglia envelopment drove myelin loss, some lost sheaths were not observed to have envelopments (**Fig. 4o-p**). Considering the longevity of sheath envelopment was frequently less than 5 h and typically less than 20 h (**Fig. 4l**), our imaging acquisition frequency (once per day) may have been insufficient to capture all envelopment events. Alternatively, some myelin loss could occur through non-phagocytic pathways. Fc gamma receptor activation can induce the release of cytotoxic factors by immune effector cells, including microglia, a process known as antibody-dependent cellular cytotoxicity, which causes apoptotic or necrotic death of target cells^58^. Downstream of complement activation, the classical cascade terminates in the formation of membrane attack complexes in the cellular membrane of targeted structures, causing cell swelling and lysis^59^. These pathways would be inhibited by the LALA Fc mutation that prevented sheath loss in this model **(Fig. 5f,i-j**), and may be responsible for the sheath blebbing phenotype we occasionally observed with MSrAb + hS treatment (**Extended Data Fig. 7b-c)**. We also occasionally observed myelin swelling (**Extended Data Fig. 7b-c**), a reversible phenotype recently described in other demyelinating models^50^. Interestingly, myelin swelling was not upregulated by MSrAb + hS treatment (Kruskal-Wallis: p = 0.28; MSrAb + hS: n = 13 mice, median (IQR) = 2% (0, 4.63) of sheaths; Saline: n = 14, median (IQR) = 0% (0, 3)) and was typically concentrated within individual oligodendrocytes (**Extended Data Fig. 7c**), suggesting swelling in this model is a disruption to some oligodendrocytes induced by surgery.

In contrast to oligodendrocyte death by laser-targeted DNA damage^60^, we did not observe microglia enveloping oligodendrocyte cell bodies, which were not targeted by antibody (**Extended Data Fig. 1b**)^11^ and remained relatively preserved (**Fig. 2e**). Without microglial or antibody targeting, we reason that the limited oligodendrocyte death we observed did not occur through microglial phagocytosis or other direct antibody-mediated mechanisms. Rather, as oligodendrocyte loss was restricted to mice with high levels of sheath loss (**Fig. 2e**) and did not occur under conditions where sheath loss was prevented (**Fig. 5e,g-h**), these cells may die in response to the overwhelming loss of their sheaths and/or through other inflammatory signaling – potentially involving microglia. Intriguingly, relative oligodendrocyte preservation is also typical in the early stages of demyelination in MS white and gray matter^61–63^ and in lesions found to contain IgG antibodies and activated complement^3,12^, while later lesion stages have almost complete loss of oligodendrocytes^61–64^. Thus, future study to elucidate the mechanisms involved in oligodendrocyte survival and death in this model may provide important strategies for protecting these cells as lesions evolve.

While microglia have been long understood to phagocytose myelin in MS lesions, we provide evidence that these cells actively drive the loss of myelin targeted by autoantibodies in MS. This process could occur in white matter lesions, which have myelin-containing microglia heavily enriched in actively demyelinating regions, including in the core of active lesions and at the expanding borders of chronic active lesions^5,38^, where we observed microglia enveloping myelin (**Fig. 4b, Extended Data Fig. 4b**). IgG antibodies and complement components associated with myelin and internalized in mononuclear phagocytes are observed in many of these lesions^3,12^, with lesions lacking these features reflecting either different lesion stages or perhaps disparities in disease mechanisms among patients^6,65^. Similarly, phagocytic destruction of myelin by microglia could occur in gray matter lesions. Cortical lesions can also harbor elevated numbers of microglia in their core or along their borders^66–69^, albeit to a lesser extent than white matter lesions^66,67,69^, and these cells phagocytose myelin^70,71^. Whether autoantibodies play a role in these lesions is currently unresolved, but some cortical lesions are found below areas of meningeal aggregates of B cells^72,70^ that are thought to release noxious factors to induce degeneration^73^ and complement opsonization of myelin in cortical lesions has been observed^69^. Importantly, BTK inhibition, which blocks microglia-mediated demyelination in our model (**Fig. 7e-i**), prevents the formation and expansion of lesions in MS white matter in clinical trials^28–31^, and may be an effective strategy to target this demyelination pathway in patients.

Beyond BTK, there may be other promising therapeutic targets along this pathway. Upstream, the protein tyrosine kinase SYK phosphorylates BTK^24^, among other targets^24,74^, and is the central mediator of Fc gamma receptor-mediated phagocytosis of antibody-bound targets by macrophages^74^. Reminiscent of our observation that inhibiting BTK preserves the envelopment of sheaths but blocks their subsequent loss (**Fig. 7**), deleting SYK preserves phagocytic cup formation but prevents the subsequent phagocytosis of the underlying structure^74^. SYK inhibition has been effective in other autoimmune diseases^75^ and may prove to be a successful strategy in MS. Downstream, BTK is involved in the important cross-talk between Fc gamma receptors and complement receptors that facilitates antibody-dependent cellular phagocytosis: following Fc gamma receptor activation, BTK activates complement receptor 3 (CR3)^24^, which aids in closing Fc gamma receptor-mediated phagocytic cups^17^. Complement-targeting therapy is a burgeoning field for the treatment of various neurodegenerative diseases, including the autoimmune demyelinating neuromyelitis optica spectrum disorder^76^. Our results indicate complement signaling is involved in inducing microglia-mediated demyelination in response to MS autoantibodies (**Fig. 1e, Fig. 5e-j**) and support targeting this pathway to prevent demyelination in MS. Moving forward, we are optimistic that continued study of microglial signaling and behavior will lead to breakthroughs in our understanding of MS pathogenesis and the development of novel therapeutic strategies for this pernicious disease.

## Author contributions

E.G.H. conceived the project. L.A.O performed experiments for Figs. 1, 2, 4, 7 and Extended Data Figs. 2, 5, 7-8 (*in vivo* imaging, optic nerve degeneration) and analyzed data for Figs. 2, 4, 7 and Extended Data Figs. 5, 7-8 (sheath tracking, morphology, and envelopment, timelapse persistence and formation of ensheathments, and code for microglia soma size and complexity). H.J.B. performed experiments for Figs. 1-4, 6-7 and Extended Data Figs. 2-3, 5, 7-8 (surgery, *in vivo* imaging, single-cell RNA sequencing). M.E.S. performed experiments for Figs. 1-4, 7 and Extended Data Figs. 3, 5, 7-8 (*in vivo* imaging, RNAscope) and analyzed data for Figs. 1, 4, 7 and Extended Data Fig. 5 (oligodendrocyte soma tracking, microglial density and envelopment status, selecting microglia for soma size and complexity analysis).

M.W. performed experiments (surgery, *in vivo* imaging) and data analysis (microglial envelopment status, oligodendrocyte soma tracking, sheath tracking and envelopment) for Fig. 5 and experiments and analysis for Fig. 4 and Extended Data Fig. 8 (CLEM). S.K. performed experiments for Figs. 1, 2 and Extended Data Figs. 2 (histology of oligodendrocyte somas, myelin, and axon degeneration) and analysis for Figs. 1, 2, 4 and Extended Data Fig. 5 (oligodendrocyte soma and myelin quantification of histology, timelapse persistence of ensheathments, microglial soma size and complexity). R.H. performed experiments for Figs. 1, 2, 4, 7 and Extended Data Figs. 5, 7-8 (surgery). G.P. performed analysis for Figs. 3, 6 and Extended Data Fig. 3 (single-cell RNA sequencing analysis).

K.S.G. performed experiments for Figs. 3, 6 and Extended Data Figs. 1, 3 (single-cell RNA sequencing, live antibody binding) and analysis for Extended Data Fig. 1 (live antibody binding). A.S. performed experiments for Figs. 4 and Extended Data Figs. 5-6 (histology of human tissue and LPC demyelination tissue). C.R.M. developed the MSrAb surgery, application, and *in vivo* imaging protocol. M.M. performed experiments and analysis for Fig. 3 and Extended Data Fig. 3 (RNAscope). D.F. performed experiments for Extended Data Fig. 8 (histology). K.H. performed experiments for Extended Data Fig. 6 (LPC surgery). C.M., C.I.T., and A.D. performed experiments and analysis for Fig. 4 and Extended Data Fig. 7 (CLEM). E.G.H. performed experiments for Figs. 1, 2, 4, and 7 and Extended Data Figs. 5, 7-8 (surgery) and analysis for Extended Data Fig. 5 (code for microglia soma size and complexity). R.G. and D.O. provided BTKi, human sequencing data, and technical expertise. G.O., J.B., and W.M. provided MSrAb and technical expertise. A.W. supervised A.S.’s and K.H.’s contributions to the project and provided expertise in MS.

E.G.H. supervised the project. H.J.B. wrote an early draft, L.A.O. wrote the current manuscript. L.A.O. made the final figures, with help from M.W., S.K., G.P., K.S.G., and A.S. L.A.O. and E.G.H. edited the manuscript with input from all other authors.

## Acknowledgements

We thank members of the Hughes, Macklin, and Bennett labs for discussions. Human microscopy and image analysis were carried out at and with the help of the IRR Imaging facility at the University of Edinburgh. LPC cryogel scaffolds were provided by Dr. Ben Newland at Cardiff University.

## Data availability

All data that support the findings, tools, and reagents will be shared on an unrestricted basis; requests should be directed to the corresponding author. The raw RNA sequencing data are available in the Gene Expression Omnibus (GEO) repository under GSE330704. The analysis pipeline for RNA sequencing and microglia morphology are available at DOI 10.5281/zenodo.20042467.

## Methods

### Animals

Animal use at the University of Colorado (all but LPC cryogels) was conducted in accordance with protocols that were approved by the Animal Care and Use Committee at the University of Colorado Anschutz Medical Campus. Mice were kept on a 14-h light–10-h dark schedule with *ad libitum* access to food and water. Male and female 25-month-old C57BL/6N *MOBP–EGFP* (MGI:4847238), *B6*.*129P2(Cg)-Cx3cr1tm1Litt/J* (Jax stock #005582), B6.129P2(Cg)-Cx3cr1tm2.1(cre/ERT2) Litt/WganJ (Jax stock #021160), and B6.Cg-Gt(ROSA)26Sortm14(CAG-tdTomato)Hze/J (Jax stock #007914) mice were used. 100 mg/kg tamoxifen was administered to *Cx3cr1-CreER; Ai14* mice by intraperitoneal injection 1-2 weeks prior to cranial window implantation. Animal use conducted at the University of Edinburgh (LPC cryogels) was carried out in accordance with the UK Home Office Animals (Scientific Procedures) Act 1986 under the project license (PP1335335), and with approval of local Bioresearch and Veterinary Services (BVS) at the University of Edinburgh. Mice were housed in ventilated cages on 12-h light–12-h dark cycle with *ad libitum* access to food and water in the BVS SCRM facility. Mice used in experiments were wild type (C57BL/6J) 8-12-week-old males.

### Generation of patient-derived recombinant antibodies

Recombinant antibodies were produced using a dual vector transient transfection system in EXPI 293 cells (ThermoFisher Scientific) and purified with protein A-sepharose (Sigma-Aldrich, St. Louis, MO), as previously described^34^. The antibody is clone MS-04-2 #30, derived from a relapsing-progressive MS patient.

### Drug treatment

A subset of mice were treated via oral gavage with PRN2675 (an analog of tolebrutinib), a CNS-penetrant, highly potent, selective, irreversible BTK inhibitor (15mg/kg, diluted in 100% PEG 200) that was provided by Sanofi. Controls were treated with vehicle (100% PEG 200). Treatment with BTKi and vehicle began 24 hours prior to surgery for MSrAb + hS application. Vehicle-treated mice were included their appropriate Saline or MSrAb + hS groups in Figs. 1, 2, and 4.

### Cranial window and application of antibodies

Young adult mice (aged 2-5 months) were anesthetized via isoflurane inhalation and kept at a temperature of 37 °C using a thermostat-controlled heating pad. The skin was cut and opened over the right cerebral hemisphere, and a 2mm × 2mm region of skull above the primary motor cortex was drilled and removed (0–2 mm anterior to bregma and 0.5–2.5 mm lateral). Under ice-cold sterile saline immersion, a cut was then made into the dura using a microbended 27.5G needle and the cortex was exposed. Sterile saline or antibody cocktails (MSrAb + hS, Mutant Fc antibody + hS, Isotype control antibody + hS, MSrAb alone, or hS alone) were bath applied to the area for a duration of 5 minutes. Then, a small cover slip (VWR, No. 1) was placed into the craniotomy and sealed into place with first Vetbond (3M) and then dental cement (C&B Metabond). Mice received a perioperative subcutaneous injection of 5 mg/kg of carprofen. To stabilize the head, a custom metal plate was fixed into place with dental cement centered above and parallel to the imaging field.

### Two photon microscopy

For *in vivo* two photon microscopy, mice were lightly anesthetized with isoflurane, kept on a thermostat-controlled heating pad, and stabilized via attachment to a custom stage. Images were taken with a Zeiss LSM 7MP microscope equipped with a BiG GaAsP detector using a mode-locked Ti:sapphire laser (Coherent Ultra) tuned to 920 nm. Stacks of a final volume of 425 μm × 425 μm × 336 μm (1,024 × 1,024 pixels) were acquired using a Zeiss W plan-apochromat ×20/1.0 NA water immersion objective. Time lapses were acquired by sampling a region of 425 μm x 425 μm x 40 μm every 45 seconds for a duration of 30 minutes. A combination of vascular and myelin landmarks were used to align imaging fields across subsequent sessions. Mice were imaged immediately post-surgery and once per day in the initial 72 hours post-surgery. A subset of mice underwent imaging two times per day, with 30 minute timelapse images. A subset of mice were re-imaged 21 days later.

### In vivo image processing and analysis

Images were analyzed using ImageJ(v1.54 f). During data analysis, experimenters were blinded to the experimental groups. Longitudinal *in vivo* two-photon 4D images were registered iteratively with Poorman3Dreg plugin for X/Y registration (ImageJv1.46r) followed by Correct3D drift plugin (EGFP channel, rigid body registration; ImageJ v1.54 f)100)^77^. For presentation in figures, image brightness and contrast levels (“Brightness/Contrast” and/or “Enhance Contrast”) were adjusted for clarity and background (“Subtract Background”) or bleed-through between channels (“Image Calculator”) was subtracted when necessary.

To conservatively track demyelination, myelin and oligodendrocytes were only referred to as “lost” if imaging resolution could be guaranteed via (1) a second channel or nearby similar object in the same channel or (2) resolution of a similar object in a deeper plane.

For oligodendrocyte cell body tracking from *in vivo* images, a custom ImageJ script^78^ enabled recording of oligodendrocyte state (new, lost, or stable EGFP+ soma) across timepoints.

For analysis of myelin sheaths from *in vivo* images, only mice with images of sufficient clarity were used. For each image, a 150 μm × 150 μm region of the highest clarity across all timepoints (at least Day 0 and Day 1, but up to Day 3) was selected. Clarity was determined by clear visibility of myelin at depth (336 μm below the surface of the brain parenchyma). Correct3D drift was then repeated on this region in layer 1: 150 μm × 150 μm × 100 μm from surface of the brain parenchyma, brightness and contrast were adjusted to optimize sheath visibility, and this ROI was used for layer 1 myelin analysis. Sheaths were traced at Day 0 using only the EGFP channel, and a custom script written by L.A.O. was used to record sheath state (“stable”, “lost”, “blebbing”, “bubbling”, “shrinking”, and whether it was enveloped by a microglia) at each following timepoint (Days 1, 2, and 3). A minimum of 50 sheaths were traced per ROI. Sheaths to be traced were selected at Day 0 from across the whole 100 μm depth, prior to visualizing their envelopment status or future loss or stability. For analysis of microglia envelopment persistence, microglia envelopment of sheaths were identified and traced at the first Day 0 timepoint immediately following surgery (0 min), and a custom script written by L.A.O. was used to record whether that sheath envelopment remained at each subsequent timepoint (30 min, 5 h, and 20 h) in time lapse images. A minimum of 25 envelopments were traced per 425 μm x 425 μm x 40 μm image. For analysis of new microglia envelopment formation, microglia envelopment of sheaths were identified and traced at 20 h, and a custom script written by L.A.O. was used to record when that sheath envelopment was present at (5 h, 30 min, or 0 min) in time lapse images. All envelopments (up to 40) were traced per 425 μm x 425 μm x 40 μm image.

For analysis of percentage of microglia envelopments that were around myelin sheaths, up to 10 layer 1 microglia with envelopments were randomly chosen per mouse from *in vivo* images from the day of surgery. The MOBP-EGFP channel was then turned on and each envelopment from the selected microglia was examined for colocalization with EGFP+ myelin. The percentage of microglial envelopments around myelin was calculated per mouse.

Given the rapid and dramatic morphological changes undergone by microglia following surgery, we could not ascertain the identity of individual microglia across time and thus performed cross-sectional analyses for density and morphology. For the percentage of microglia with envelopments per day (Days 0, 1, and 2), all layer 1 microglia with at least one envelopment were counted and divided by the total microglial number at each respective timepoint. For microglia density, all layer 1 microglia per 425 μm × 425 μm × 100 μm ROI were counted using Cell Counter (Fiji) on Day 0 and Day 2. For microglia morphology, 3 layer 1 microglia were selected per *in vivo* image from both Day 0 and Day 2 timepoints. Selected microglia were those where the entirety of the cell body and processes were visible and distinguishable from adjacent microglia. The “Selection Brush Tool” was used to select all portions of the microglia in each z plane, while the other image data was cleared. A custom script written by E.G.H. and edited by L.A.O. and S.K. (available at DOI 10.5281/zenodo.20042467) was used to process the image to derive the soma region and skeletonize and quantify the processes. To summarize, soma filtering was done with Gaussian Blur and the 3D Filters plugin (Minimum and Maximum)^79^. We noticed that due to reduced clarity in some cranial windows, some images were acquired with higher laser power and required different filtering settings to accurately derive the soma size. Thus, two different filtering parameters were used, with all images passing through the first filtering settings, and the alternative filtering settings only used if the first filtering settings did not accurately outline the soma as determined by user judgement. The bounds of the soma were then detected using the 3D Objects Counter plugin^80^ and the area quantified. The few microglia whose properties were not accurately captured by these settings were manually traced to define the soma. Filtering of microglia processes was done using the same settings for all images. Images were filtered with Maximum Filter, Mean Filter, Despeckle, and Subtract Background. Images were then Autothresholded, Despeckled again, and outliers were removed (Remove Outliers). The processed images were then Maximum Intensity Z Projected and Skeletonized. The area of the soma was removed, and the skeletonized process complexity was analyzed with a Sholl Analysis.

### Single-cell RNA sequencing and analysis

For scRNAseq, micro-dissection of cortical tissue (from brain surface to corpus callosum) was performed from a 2 × 2 mm area beneath site of MSrAb + hS application and from the intact contralateral cortex 72 h post-application. Tissue was pooled from 3 mice expressing GFP in cortical microglia (*Cx3Cr1-EGFP*) per condition. Tissue was dissected, minced and then dissociated with Miltenyi’s Neural Dissociation Kit (P) following manufacturer’s instructions. Tissue was incubated in Enzyme Mix 1 for 15m at 37 °C, Enzyme Mix 2 was added, and sample chunks were gently pipetted with a P1000. Samples were incubated 10 additional minutes at 37 °C and remaining cells dissociated by gently pipetting with a flame polished glass pipette. Cells were filtered through a 40 μm filter and pelleted at 400 × g for 10 min to remove enzyme. Cells were resuspended and debris was removed using Miltenyi Debris Removal Solution. Remaining cells were FACS sorted for singlet, live (DAPI-negative), EGFP+ cells.

Sorted cells were captured using the 10X Chromium V3 platform and sequenced to approximately 60,000 reads per cell using the Illumina NovaSEQ 6000. Sample demultiplexing, barcode processing, and alignment was performed using Cell Ranger (10x Genomics). Processed data was analyzed in R using Seurat version 5.1.0^81^. For initial quality control, cells containing fewer than 1000 detected genes and more than 7% percent mitochondria were excluded from subsequent analysis. Cells containing more than 10,000 UMIs were excluded. The contribution of the percentage mitochondrial and ribosomal gene reads to variance between cells was regressed out during data normalization and scaling. Unsupervised clustering was performed using a resolution of 0.5 in a UMAP embedding space also used for visualization. MSrAb + hS samples from Fig. 3 were reanalyzed together with BTKi + MSrAb + hS samples in Fig. 6. To annotate clusters, the differentially expressed gene lists resulting from the Wilcoxon rank sum test were manually inspected, and names assigned based on gene enrichment per cluster. One-vs-all cluster marker analysis is presented in **Extended Data Table 2**. Differential expression analysis across clusters was performed using multiple comparisons adjusted Wilcoxon rank sum test. We used an absolute fold-change over 2 and an adjusted p-value below 0.01 to define a meaningful change in gene expression. Pathway analysis was performed using the SCPA package using Gene Ontology Biological Process term lists^82^. The raw data are available in the Gene Expression Omnibus (GEO) repository under GSE330704. The analysis pipeline is available at DOI 10.5281/zenodo.20042467.

### Human RNA sequencing and analysis

These data were derived from a previously published dataset^26^. A Student’s t test was performed on selected genes to determine differential expression between groups. Genes were considered significantly altered if the absolute fold-change in expression was ≥ 1.2 and the p-value < 0.05.

Briefly, frozen human brain tissues were obtained from the Human Brain and Spinal Fluid Resource Center at University of California Los Angeles. Post-mortem derived tissue (50 mg) was homogenized and lysed in 1% Triton lysis buffer (TBS). After centrifugation (13,000 × g), supernatant was collected as the soluble fraction. Lysis buffers contained protease and phosphatase inhibitors (1:100 dilution; Thermo Fisher Scientific, #78444). For bulk RNA library preparation and sequencing, RNA was extracted from 25 frozen brain sections according to the “Purification of Total RNA from Microdissected Cryosections” method of the Qiagen RNeasyMicro kit, with DNase I (Cat# 74004; Qiagen). Sections were maintained separately throughout processing. RNA integrity and quantification were determined using a Bioanalyzer RNA 6000Pico chip on the Bioanalyzer 2100 (Cat# 5067-1513, Agilent). Strand-specific RNASeq libraries were created using the TruSeq stranded mRNA library prep Kit (cat# RS-122-2101, Illumina). The library sizes and quantification were determined using the Bioanalyzer High Sensitivity DNA kit (Cat# 5067-4626, Agilent). The libraries were sequenced using the HiSeq2000 (Illumina). All data processing from the QC of FASTQ files to the derivation of DEGs was performed within Array Studio (Version 10,Omicsoft Corporation, Research Triangle Park, NC, USA). Data quality was assessed using the “Raw Data QC Wizard” function within ArrayStudio. The sequence used to trim the adapters from the reads was “AGATCGGAAGAGCG.” Paired reads were mapped to the human reference genome (GRCh38, GenCode.V24) using the Omicsoft Aligner4 (OSA4)^83^. The Expectation-Maximization algorithm was implemented to calculate the FPKM (Fragments Per Kilobase Million) value for each gene^84^. Lowly expressed genes were filtered out. Additionally, the genes were filtered to retain only protein coding genes.

### RNAscope and analysis

For RNAscope, in situ hybridization (ISH) was performed using the RNAscope Multiplex Fluorescent Reagent Kit V2 (ACD Biotechne 323100). Probes were designed for mouse Ccl4-C1 (ACD Biotechne 421071), Fcrls-C2 (ACD Biotechne 441231) and Cd72-C3 (ACD Biotechne 1046761). 20 μm paraformaldehyde-fixed brain sections were washed and mounted on Superfrost microscope slides. Tissue dehydration, antigen retrieval, protease treatment, probe hybridization, amplification, and horseradish peroxidase (HRP) reaction steps for C2, C3 and C1 (in this order) were performed according to the manufacturer’s protocol. All washes were for 2 min in 1X RNAscope wash buffer at room temperature unless otherwise noted. For amplification, slides were washed 2x before incubation with Amp 1 for 30 min at 40 °C. This process was then repeated for Amp 2 and Amp 3. For the HRP reaction, slides were then washed 2x before incubation with the respective HRP conjugate for 15 min at 40 °C. Next, sections were washed 2x and incubated in the corresponding fluorophore for 30 min at 40 °C (C2, Tyramide Signal Amplification (TSA) Vivid 570 1:1,500 in TSA buffer, C3 TSA Vivid 650 1:2,000 in TSA buffer, or C1 Opal TSA-digoxygenin (Akoya) 1:750 in TSA buffer). HRP reaction slides were then washed 2x, incubated in HRP blocker for 15 min at 40 °C and finally washed 2x. This was then repeated for the remaining two probes. Following the final HRP blockade, slides were incubated in blocking buffer (5% normal donkey serum and 0.3% Triton X-100 in PBS) for 60 min at room temperature. Slides were then incubated with chicken anti-GFP 1:250 (AVES, GFP-1020) for 48 h at 4 °C. Slides were next washed 3x for 15 min in PBS. Immunostaining was completed with incubation in Donkey anti-chicken 488 1:500 (Jackson, 703-545-155) for 60 min. Note that GFP immunostaining was not analyzed due to technical difficulties. ISH was completed by washing slides 3x for 15 min in PBS before incubation in Polaris 780 (1:125 in diluent buffer, Akoya) for 30 min at 40 °C. Finally, slides were washed 2x and counterstained with RNAscope 4,6-diamidino-2-phenylindole (DAPI) for 30 s before cover slipping with Prolong Gold mounting medium (Invitrogen P36934) for imaging.

Tiled fluorescent images of cortex were acquired using the Olympus VS200 microscope with a 40X air objective and a 10% tile overlap. A *z*-stack with 0.84 μm steps captured the entire thickness of each tissue section. Images were stitched and maximum intensity projected before analysis.

Images were analyzed using Qupath Software 0.4.3. RNA expression per cell was quantified using a custom analysis pipeline. An ROI for cortical layer 1 was manually drawn onto each tissue section.

Nuclei were semi-automatically segmented based on DAPI. Nuclear segmentations were dilated to encompass the predicted area of each cell. Finally, mRNA puncta were segmented and assigned to DAPI+ cell regions. Threshold values for each channel were based on the background signal in each image. Cell and puncta detection was performed on each section in Qupath and the spreadsheets containing the intensity and number of puncta within each cell were exported and processed. Finally, the percentage of *Ccl4*+*Fcrls*+/*Fcrls*+ and *Cd72*+*Fcrls*+/*Fcrls*+ cells were calculated using the established thresholds.

### Live antibody binding assay

Sagittal cerebellar slices (300 μm) were prepared from P10 *PLP-EGFP* mice and cultured on MilliCell 0.4 μm membrane inserts (Millipore, Billerica, MA) for 7-10 days in slice media (25% Hank’s balanced salt solution (HBSS), 25% heat-inactivated horse serum, 50% minimum essential media (MEM), 125 mM HEPES, 28 mM D-Glucose, 2 mM L-Glutamine, 10 U/ml penicillin/streptomycin, all from Life Technologies, Carlsbad, CA) at 37 °C. For live binding, unfixed slices were incubated with 20 μg/ml recombinant antibodies for 1 h at 37 °C. Slices were then rinsed 1x in PBS and fixed in 4% PFA for 20 min.

### Optic nerve degeneration

To generate a positive and negative control for neurodegeneration immunostaining, mice were euthanized with an intraperitoneal injection of sodium pentobarbital (100 mg/kg). Both optic nerves were immediately dissected, and placed on a laminated paper to remain flat during processing. One optic nerve was immediately post-fixed in 4% PFA on ice for 1 h (negative control). The other optic nerve was incubated in DMEM/F-12 at 37 °C for 6 h to allow for degeneration to proceed, then post-fixed in the same manner as the first optic nerve. Following fixation, optic nerves were stored in 30% sucrose at 4 °C until sectioning.

### Cryogel implantation

Cryogel scaffolds were produced by Dr. Benjamin Newland and Dr. Sophie Hill at Cardiff University. The cryogels are 2 mm diameter × mm depth cylindrical sponge-like scaffolds, made from polyethylene glycol^85^. Cryogels were placed in 250 μl of 10mg/ml L-α-Lysophosphatidylcholine (LPC, L4129, Sigma-Aldrich, batch SLCM5569) or sterile PBS. Craniotomies were performed in anesthetized 8-12-week-old wild-type male mice, by drilling a 2 mm diameter hole over the right primary motor cortex, localized relative to Bregma. As previously described^49^, a LPC or PBS-filled cryogel was placed onto the cortical surface and the skin was sutured closed over the cryogel.

### Tissue processing, immunostaining, imaging, and analysis

Mice were euthanized with an intraperitoneal injection of sodium pentobarbital (100 mg/kg) and transcardially perfused with ice-cold 0.1M phosphate buffered saline (PBS) and 4% paraformaldehyde. Brains were post-fixed with 4% paraformaldehyde overnight, then stored in 30% sucrose at 4 °C for at least 24 hours. Brains were sectioned coronally at either 10 or 30 μm thickness using a cryostat. Only the right (surgerized) hemisphere was sectioned. Optic nerves were sectioned longitudinally at 10 μm thickness. Immunostaining was done on free-floating sections (brain) or mounted section (optic nerve). Sections were incubated in blocking solution (5% donkey or 10% goat serum, 0.2% Triton X-100 in PBS) at room temperature for 2 hours, then incubated with primary antibody overnight at 4 °C. Sections were washed with blocking solution and then incubated with secondary antibodies for 2 hours at room temperature. Sections were mounted on slides using Vectashield antifade mounting media. Immunostained brain sections were imaged with a laser-scanning confocal microscope (Nikon A1R). All images were quantified with the user blinded to condition.

For oligodendrocyte and myelin analysis, 3 coronal sections from Bregma +1.1 mm (under the cranial window) were used for each mouse. 10X tiled z stack images were acquired spanning the dorsal and dorsal-lateral layer 1-3 of cortex. The same region was quantified for each mouse. EGFP+ oligodendrocytes were quantified in ImageJ(v1.54 f) from Maximum Intensity Z Projected images. To quantify myelin, machine learning-based bioimage analysis software ilastik^86^ was used. For each image, the software was trained by a skilled user on positive structures (MBP+ myelin) and negative structures (background). ilastik then derived the positive and negative pixels for the entire image, which were then measured in ImageJ(v1.54 f).

For immunostaining of cerebellar slices, slices were rinsed in PBS, blocked and permeabilized for 30 min (5% NDS, 1% Triton X-100 in PBS) and incubated in primary antibody overnight at room temperature with agitation. Slices were washed 3 × 10 min in PBS, then secondary antibody was applied for 2 h at room temperature. Slices were washed again 3 × 10 min in PBS and finally mounted onto slides with Fluoromount G. Confocal images were acquired by Leica SP5 laser scanning microscope (Leica Microsystems GmbH, Wetzlar, Germany). Colocalization of recombinant antibodies and PLP-EGFP was automated using Cell Profiler. Single channels were thresholded and the area of positive pixels was measured. The recombinant antibody positive pixel area was swelled by 1 pixel (expanded) and masked against thresholded PLP-EGFP. % PLP-EGFP colocalized with recombinant antibody = masked area/total PLP-EGFP area.

For immunostaining of LPC cryogel mice, mice were anesthetized by an intraperitoneal injection of ketamine and medetomidine 1-week post-surgery during the early demyelination phase and euthanized by transcardially perfusion with 4% paraformaldehyde in PBS (v/v). Brains were post-fixed with 4% PFA for 18-24 h at 4°C. Samples were stored in increasing concentrations of sucrose (15%, 30%) and were subsequently chemically frozen in 2-methyl-butane to be stored at −80°C. Brains were coated in CryomatrixTM embedding resin (Epredia, 6769006) and cryosectioned into 10 μm sections over the lesion area. Slides were submerged in 70% ethanol in a coplin jar at −20 °C for 15 min. Citric acid-based antigen retrieval was performed by microwaving sections for 15 min in Antigen Unmasking Solution (Vector Laboratories, H-3300) and cooled for 10 minutes in room temperature tap water. Sections were blocked for 1 h using 10% normal horse serum in 0.5% Triton X-100 in Tris-buffered saline. Incubation with primary antibodies occurred overnight at 4 °C under humidified conditions. The next day, sections were incubated for 1 h at room temperature with Alexa Fluor secondary antibodies and counterstained with Hoechst to visualize nuclei. Slides were mounted with SouthernBiotech Fluoromount-G slide mounting medium (Cambridge Bioscience, 0100-01). Whole sections were imaged using an Opera Phenix Plus with a 20x air objective (Zeiss, NA 0.4), with increased numbers of microglia in L1-2/3 of the right cortex indicating the early demyelinating region. High-resolution images were obtained using a Leica SP8 4D confocal microscope with a 63X objective (Harmonized Compensation Plan Apochromatic CS2 63x oil, NA 1.4). Z-stacks were collected using an experiment-specific optimal z-step.

For a full list of antibodies see **Extended Data Table 7**.

### Human tissue processing, immunostaining, and imaging

Human formalin-fixed paraffin-embedded (FFPE) brain tissue sections from MS patients and controls were provided by a UK prospective donor scheme with full ethical approval from the UK Multiple Sclerosis Society Tissue Bank (MREC/02/2/39) and from the MRC-Edinburgh Brain Bank (16/ES/0084). Using neuropathological methods, MS diagnosis with MS was confirmed by F. Roncaroli (Imperial College London) and C. Smith (Centre for Clinical Brain Sciences, Centre for Comparative Pathology, Edinburgh) with no indication of confounding neurodegenerative diseases. Clinical history was provided by R. Nicholas (Imperial College London) and C. Smith. FFPE tissue blocks were sliced into 4-μm sequential sections of 2 cm x 2 cm x 1 cm and stored at room temperature. All samples chosen were from cortical gray and white matter, and demyelination was confirmed in MS samples. For human patient metadata see **Extended Data Table 5**.

FFPE sections were dewaxed and rehydrated using xylene and decreasing concentrations of ethanol. Citric acid-based antigen retrieval was performed by microwaving sections for 15 minutes in Antigen Unmasking Solution (Vector Laboratories, H-3300) and cooled for 20 minutes in room temperature tap water. Tissue autofluorescence was quenched with Autofluorescence Eliminator Reagent (Merck, 2160), sections were washed with washing buffer (0.001% Triton X-100, Tris-buffered saline) and subsequently with 3% H_2_O_2_. Sections were blocked for 1 h using 10% normal horse serum in 0.5% Triton X-100 in Tris-buffered saline. Incubation with primary antibodies occurred overnight at 4°C under humidified conditions. The next day, sections were incubated for 1 h at room temperature with Alexa Fluor secondary antibodies and counterstained with Hoechst to visualize nuclei. Slides were mounted with SouthernBiotech Fluoromount-G slide mounting medium (Cambridge Bioscience, 0100-01). For a full list of antibodies see **Extended Data Table 7**.

Whole sections were imaged using an Opera Phenix Plus with a 20x air objective (Zeiss, NA 0.4), with a lack of PLP1 staining being used to identify lesions. High-resolution images were obtained using a Leica SP8 4D confocal microscope with a 63X objective (Harmonized Compensation Plan Apochromatic CS2 63x oil, NA 1.4). Z-stacks were collected using an experiment-specific optimal z-step.

### Correlative light electron microscopy

On Day 1, live *in vivo* two-photon imaging was performed, as described above, in a region of interest containing several microglial myelin envelopments. Immediately following imaging, the mouse was euthanized with an intraperitoneal injection of sodium pentobarbital (100 mg/kg) and transcardially perfused at a flow rate of 1.6 mL/min with ice-cold 0.9% NaCl containing 2mM CaCl_2_ followed by fixative consisting of 2.5% glutaraldehyde and 2% paraformaldehyde in 0.1 M phosphate buffer. Post-perfusion, the region of interest was identified on the two-photon microscope and was delineated by performing near infrared branding (NIRB) using the two-photon laser tuned to 780nm. The brain was then dissected and post-fixed in 2.5% glutaraldehyde and 2% paraformaldehyde in 0.1 M phosphate buffer for 2 h at room temperature, then stored at 4°C in 0.1 M phosphate buffer. The brain was subsequently trimmed down and embedded in 4% low melt-point agarose. A 200 μm horizontal section of the branded area was collected using a vibratome and re-imaged on the two-photon microscope at 2048 × 2048 resolution to enable alignment of fluorescence and EM images. The sample was then processed for electron microscopy, and the region of interest was identified as previously described^87^. Following resin embedding, serial ultrathin sections (70 nm) were cut using an ultramicrotome (UC Enuity, Leica Microsystems). Initial sections were collected on copper grids and examined using a JEM-120i transmission electron microscopy (JEOL) to assess tissue preservation and verify the location of the region of interest by comparing to the location of burn marks and blood vessels to the two-photon image of the horizontal section. Consecutive serial sections were then collected manually onto silicon wafers and imaged using a Gemini 460 scanning electron microscope (Zeiss) equipped with a volumeBSD backscatter detector. Final alignment of structures was performed by comparing the location of blood vessels and myelin sheaths between two-photon and EM images.

### Statistics

Sample sizes in this study were not predetermined statistically, but were comparable to relevant publications. Due to the extremely low throughput of these experiments, in Fig. 1, 2, and 4, mice receiving vehicle treatments (for BTKi) were also included with their respective groups (Saline or MSrAb + hS). Statistical analyses were conducted using JMP17 or 19 (SAS), Prism 7 (GraphPad), or R. Two-tailed tests and α < 0.05 were always employed unless otherwise specified. When normality or equal variance was not assumed (e.g. oligodendrocyte or sheath loss in Saline versus MSrAb + hS group from *in vivo* images), we used nonparametric tests, namely Kruskal-Wallis with Dunn post-hoc comparisons (if applicable). For normally-distributed data, we used parametric statistics, including paired or unpaired two-tailed Student’s t-tests, or analysis of variance (ANOVA) with Tukey’s honestly significant difference (HSD). Analysis of survival curves was done with a Log-Rank test. Paired tests were always within mouse. For data visualization, nonparametric data is presented with median and IQR, while parametric data is presented with mean and standard error of the mean. Full statistical details are presented in **Extended Data Table 1**.

