## Supplemental Figures for "Microglia drive demyelination via multiple sclerosis antibodies and BTK signaling"

[1]Department of Cell and Developmental Biology, [2]Department of Physiology and Biophysics, [3]Department of Neurology, [4]Department of Ophthalmology, University of Colorado School of Medicine, Aurora, CO, USA, [5]Centre for Regenerative Medicine, Institute for Regeneration and Repair, MS Society Edinburgh Centre for MS Research, The University of Edinburgh, Edinburgh, UK, [6]Neurology Research, Sanofi, Framingham, MA, USA

*These authors contributed equally.

**Extended Data Video and Table Legends**

**Extended Data Video 1: IBA1+ cell-myelin interaction through z in MS chronic active WM lesion border.** 63X confocal z stack from Fig. 4C of IBA1+ cell (magenta) enveloping a myelin sheath (white) in an MS chronic active WM lesion border**.** Z slice indicated in upper left. Scale bar is 10 $\mu$m.

**Extended Data Video 2: IBA1+ cell-myelin interaction through z in MS chronic active WM lesion border.** 63X confocal z stack from Extended Data Fig. 4B (left) of IBA1+ cell (magenta) enveloping a myelin sheath (white) in an MS chronic active WM lesion border. Z slice indicated in upper left. Scale bar is 5 $\mu$m.

**Extended Data Video 3: IBA1+ cell-myelin interaction through z in MS chronic active WM lesion border.** 63X confocal z stack from Extended Data Fig. 4B (right) of IBA1+ cell (magenta) enveloping a myelin sheath (white) in an MS chronic active WM lesion border. Z slice indicated in upper left. Scale bar is 5 $\mu$m.

**Extended Data Video 4: Dynamics of microglial envelopments.** Microglia envelopments (magenta, termini indicated by white arrows) around myelin (white) can be observed growing and or shrinking over an *in vivo* 30 min timelapse. Scale bar is 20 $\mu$m.

**Extended Data Video 5: Microglia envelopment precedes sheath loss.** Two microglia envelopments (magenta, termini indicated by white arrows) around myelin (white) can be observed growing to cover the entirety of a myelin sheath over 30 min *in vivo*. By 20 h, the sheath is lost. Scale bar is 20 $\mu$m.

**Extended Data Table 1: Statistical details**

**Extended Data Table 2: Genes defining clusters**

**Extended Data Table 3: Differential gene expression between clusters**

**Extended Data Table 4: Differential gene expression between groups**

**Extended Data Table 5: Human patient metadata**

**Extended Data Table 6: Gene expression changes with BTK inhibition**

**Extended Data Table 7: Antibodies for immunostaining**

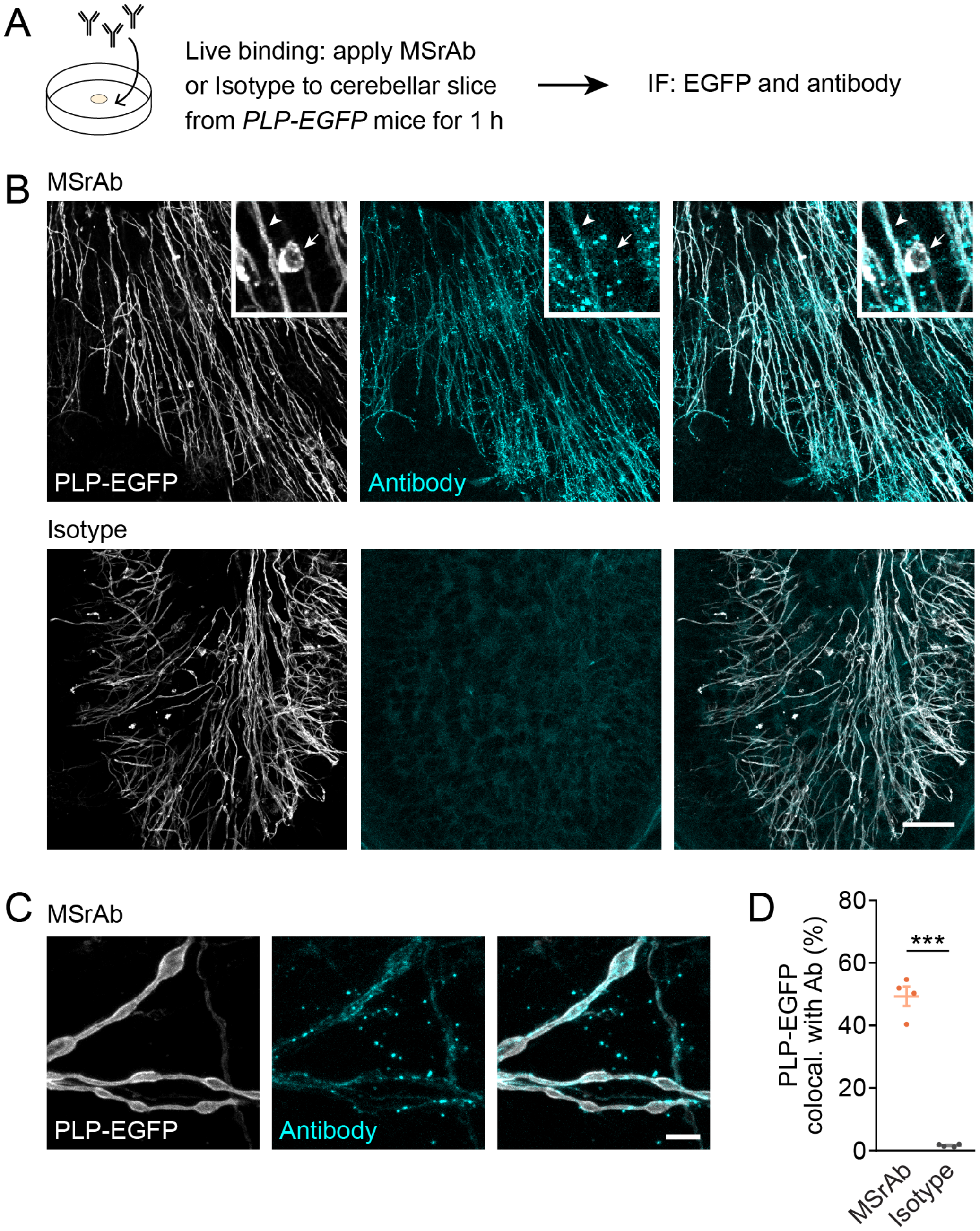


**Extended Data Fig. 1: MS patient-derived recombinant antibodies bind to myelin. (A)** Experimental protocol for antibody live binding assay and immunostaining of *PLP-EGFP* cerebellar slices. **(B)** Representative images of cerebellar slices showing colocalization of MSrAb but not an Isotype antibody (cyan) with EGFP+ myelin (white). Scale bar is 50 $\mu$m. Inset shows MSrAb colocalization with EGFP+ myelin (arrowhead) but not EGFP+ oligodendrocyte somas (arrow). **(C)** Higher magnification (60X objective) images of MSrAb (cyan) colocalization with EGFP+ myelin (white). Scale bar is 5 $\mu$m. **(D)** Quantification of % of EGFP colocalized with antibodies reveals high levels of binding of MSrAb to myelin, but no myelin binding by an Isotype antibody. t test ($t(6) = 15.3$, $p < 0.0001$; MSrAb: $n = 4$, mean $\pm$ SEM = $49.33 \pm 3.11$; Isotype: $n = 4$, mean $\pm$ SEM = $1.644 \pm 0.2171$). $p < 0.05$, ** $p < 0.01$, *** $p < 0.001$, n = slices; two-sided statistical test. See **Extended Data Table 1** for statistical details. IF = immunofluorescence.

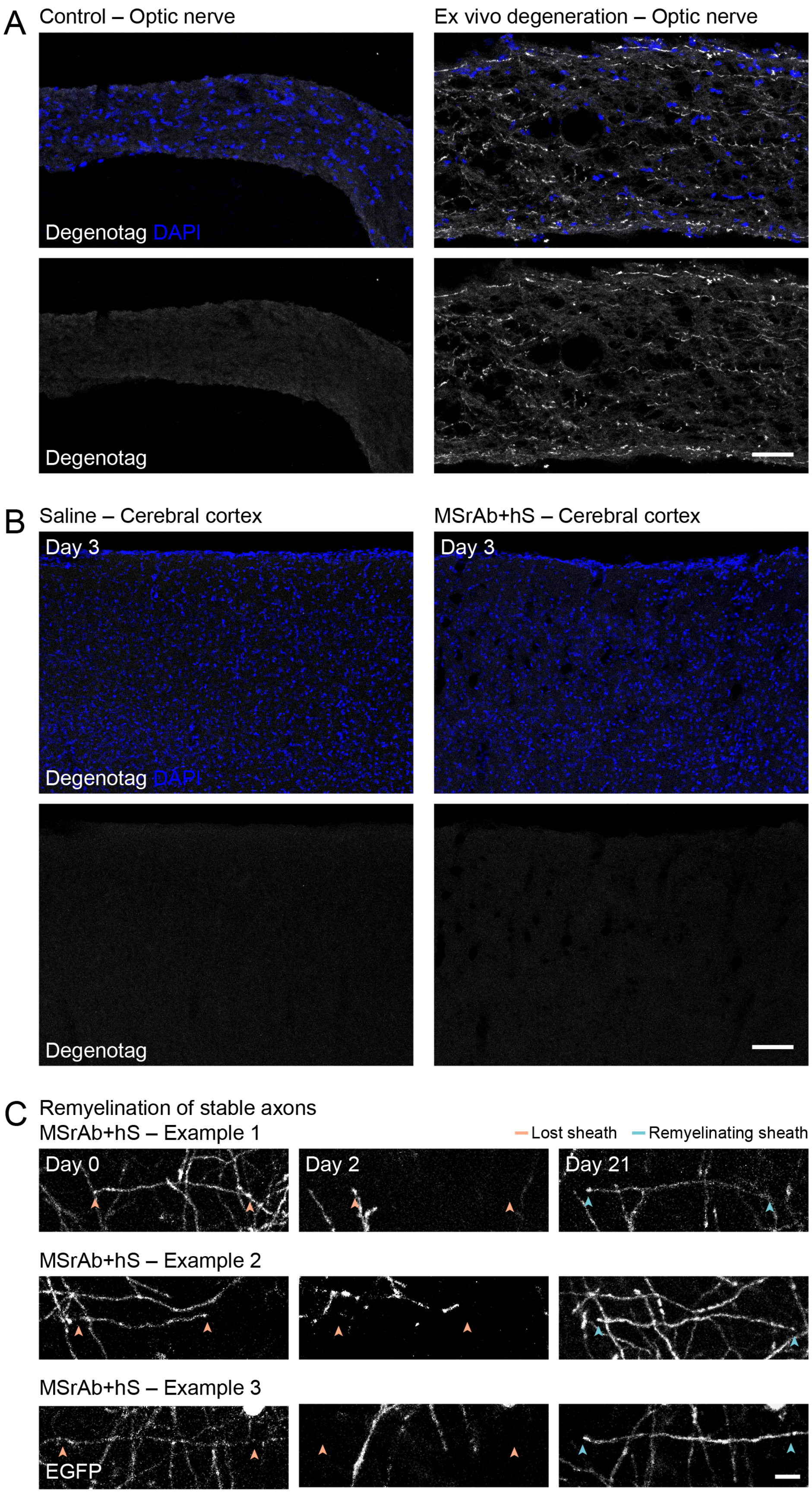
A
Control – Optic nerve
Ex vivo degeneration – Optic nerve
Degenotag DAPI
Degenotag
B
Saline – Cerebral cortex
MSrAb+hS – Cerebral cortex
Day 3
Day 3
Degenotag DAPI
Degenotag
C
Remyelination of stable axons
MSrAb+hS – Example 1
Lost sheath
Remyelinating sheath
Day 0
Day 2
Day 21
MSrAb+hS – Example 2
MSrAb+hS – Example 3
EGFP

**Extended Data Fig. 2: No evidence of axon degeneration in MSrAb demyelination model. (A)** Representative images of negative and positive control tissue demonstrating antibody specificity to degeneration. Degenerating axons are clearly labeled in degenerating mouse optic nerve tissue (right). Scale bar is 50 $\mu$m. **(B)** Representative images of Saline and MSrAb + hS cortical tissue shows no staining for degeneration on Day 3. Scale bar is 100 $\mu$m. **(C)** Example *in vivo* images of remyelination (by Day 21) of axons that were demyelinated over the first two days following MSrAb + hS application, evidencing their stability through the demyelinating insult. Scale bar is 10 $\mu$m.

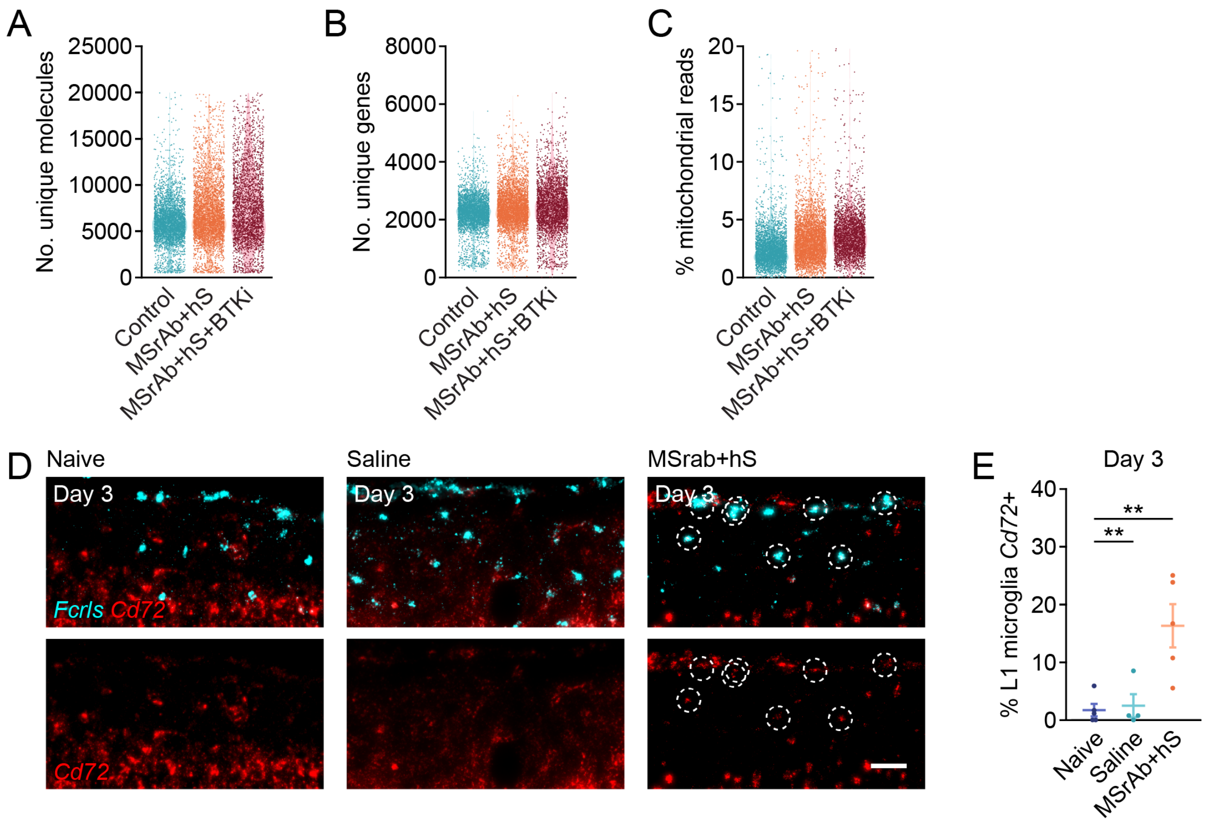


**Extended Data Fig. 3: Single-cell RNA sequencing quality control and validation. (A-C)** Number of unique molecules (A), number of unique genes (B), and percent of reads mapped to mitochondrial genes (C) in individual cells in Control, MSrAb + hS, and MSrAb + hS + BTKi samples. **(D)** Representative images of RNAscope on Day 3 reveals upregulation of *Cd72* in layer 1 microglia (*Fcrls*+) in MSrAb + hS treated mice. *Cd72+Fcrls*+ microglia are outlined with a dashed circle. Scale bar is 50 μm. **(E)** Increase in % of layer 1 microglia expressing *Cd72* on Day 3 by RNAscope in MSrAb + hS treated tissue. ANOVA ($F$ (2, 11) = 10.2492, $p = 0.0031$; Naïve: $n = 5$, mean ± SEM = 1.7345 ± 2.529; Saline: $n = 4$, mean ± SEM = 2.4947 ± 2.827; MSrAb + hS: $n = 5$, mean ± SEM = 16.3390 ± 2.529. Post-hoc Tukey's: MsrAb + hS vs. Naive ($p = 0.0047$), MsrAb + hS vs. Saline ($p = 0.0098$). * $p < 0.05$, ** $p < 0.01$, *** $p < 0.001$, n = mice; two-sided statistical tests. See **Extended Data Table 1** for statistical details.

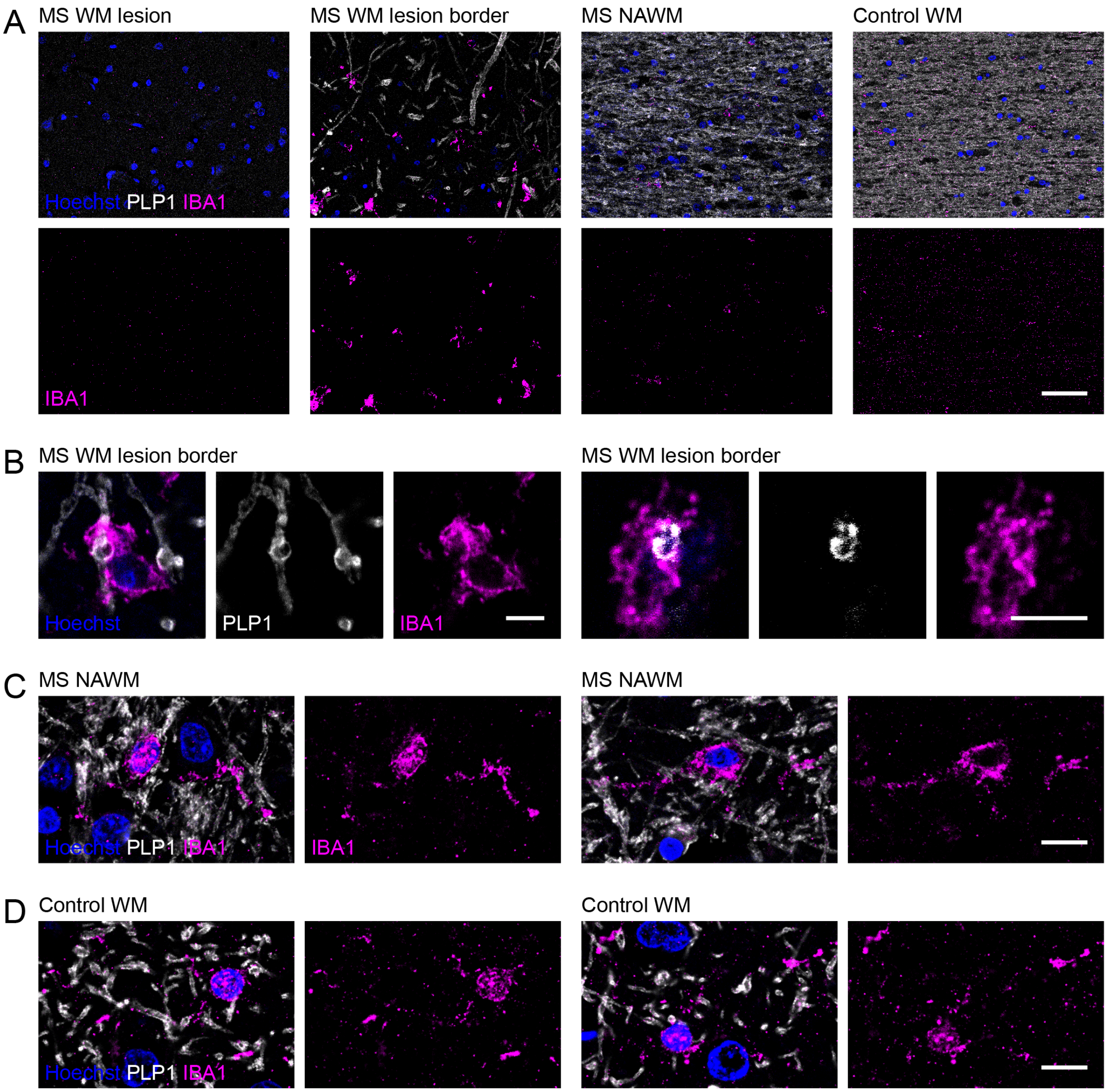


**Extended Data Fig. 4: IBA1+ cell-myelin interactions in MS chronic active WM lesion border but not NAWM or control WM. (A)** Representative images of IBA1+ cell density (magenta) shows enrichment and activation of IBA1+ cells in MS chronic active WM lesion border as compared to a chronic active WM lesion or NAWM from the same patient or WM from a control individual. Myelin staining (white) was used to defined lesion, lesion border, NAWM, and control WM. Scale bar is 50 μm. **(B)** Example 63X confocal images of IBA1+ cells (magenta) enveloping myelin sheaths (white) in an MS chronic active WM lesion border. Scale bars are 5 μm. See **Extended Data Videos 2 and 3** to view these microglia-myelin interactions through z. **(C)** No clear interactions of IBA1+ cells (magenta) and myelin (white) in NAWM in same patient. Scale bar is 10 μm. **(D)** No clear interactions of IBA1+ cells (magenta) and myelin (white) in control WM. Scale bar is 10 μm. WM = white matter; NAWM = normal-appearing white matter. See **Extended Data Table 5** for human patient metadata.

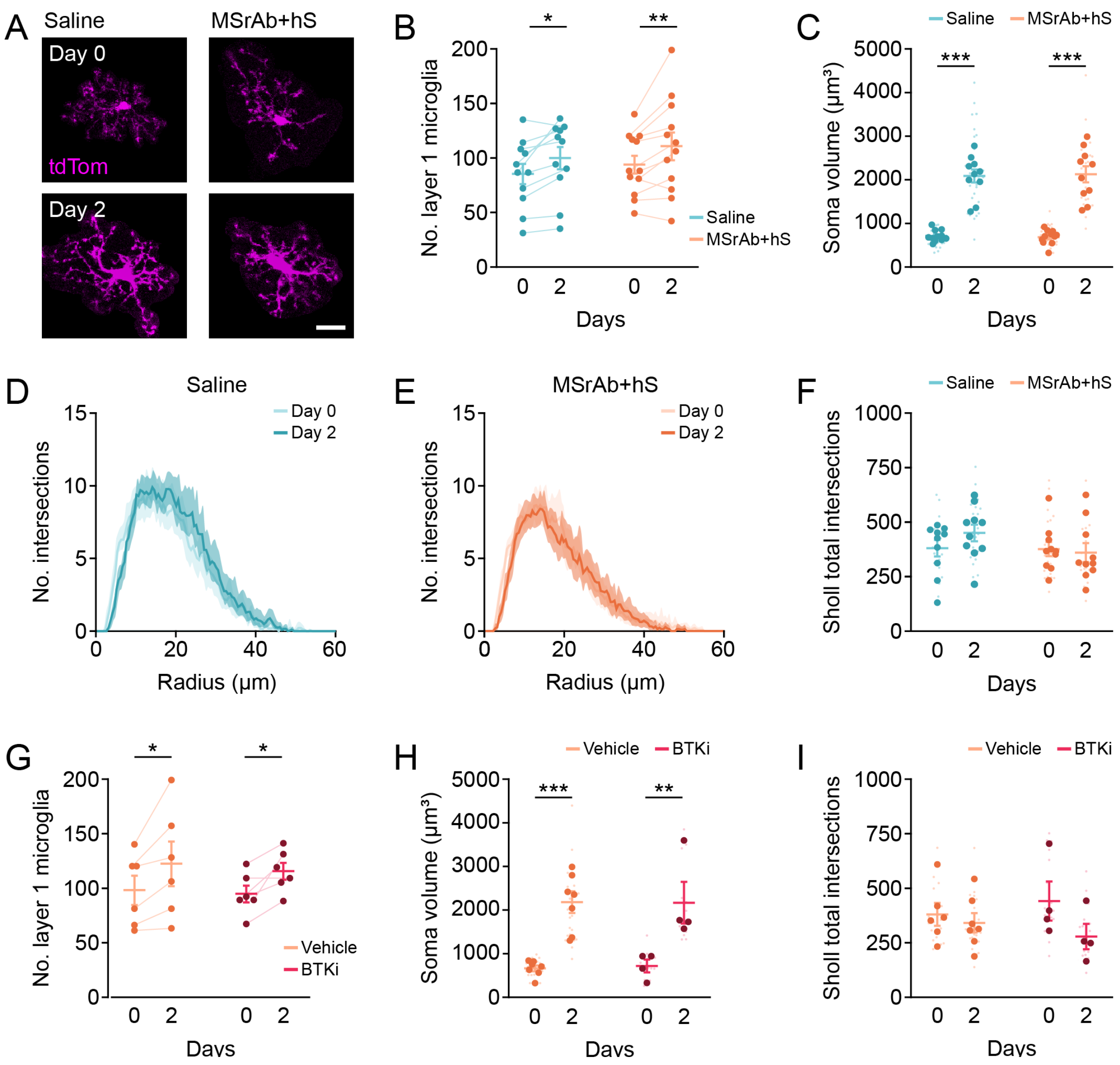


**Extended Data Fig. 5: Response of microglia density and morphology to surgery but not MS patient-derived recombinant antibodies or BTKi. (A)** Representative *in vivo* images of individual layer 1 microglia in Saline and MSrAb + hS conditions showing an increase in soma size in both groups from Day 0 to Day 2. Scale bar is 20 μm. **(B)** Microglia number is increased from Day 0 to Day 2 in Saline and MSrAb + hS treated mice. **(C)** Microglia soma volume is increased from Day 0 to Day 2 in both conditions. **(D-F)** Sholl analysis plots for microglia process complexity in Saline (D) and MSrAb + hS (E) treated mice at Day 0 and Day 2. Data shown as moving average with 95 % CI. N as in **F**. **(F)** No differences in microglia complexity as measured by Sholl total interactions over time or between groups. **(G)** BTKi inhibition does not impact surgery-induced microglia density increase. **(H)** BTK inhibition does not impact surgery-induced microglia soma volume increase. **(I)** No differences in microglia complexity as measured by Sholl total interactions over time or between groups. In **B**, Repeated measures ANOVA: Group ($F(1, 21) = 0.4753$, $p = 0.4981$), Time ($F(1, 21) = 17.5692$, $p = 0.0004$), Interaction ($F(1, 21) = 0.0934$, $p = 0.7629$). Saline Day 0: n = 11, mean ± SEM = 85.2727 ± 9.3255. Saline Day 2: n = 11, mean ± SEM = 99.8182 ± 10.2308. MSrAb + hS Day 0: n = 12, mean ± SEM = 93.8333 ± 8.1071. MSrAb + hS Day 2: n = 12, mean ± SEM = 110.6667 ± 12.7281. Post-hoc t test with Bonferroni correction: Saline Day 0 vs. Day 2 ($F(21) = 2.6899$, $p = 0.0274$),

MSrAb + hS Day 0 vs. Day 2 (F (21) = 3.2515, p = 0.0076). In **C**, Two-way ANOVA: Group (F (1, 38) = 0.0034, p = 0.9535), Time (F (1, 38) = 142.2, p < 0.0001), Interaction (F (1, 38) = 0.0821, p = 0.7760). Saline Day 0: n = 11, mean ± SEM = 701.4633 ± 39.3735. Saline Day 2: n = 10, mean ± SEM = 2084.7752 ± 150.5081. MSrAb + hS Day 0: n = 11, mean ± SEM = 674.3810 ± 50.3062. MSrAb + hS Day 2: n = 10, mean ± SEM = 2125.8004 ± 186.6351. Post-hoc Bonferroni's: Saline Day 0 vs. Day 2 (F (38) = 8.231, p < 0.0001), MSrAb + hS Day 0 vs. Day 2 (F (38) = 8.636, p < 0.0001). In **F**, Two-way ANOVA: Group (F (1, 36) = 1.513, p = 0.2266), Time (F (1, 36) = 0.483, p = 0.4915), Interaction (F (1, 36) = 1.232, p = 0.2744). Saline Day 0: n = 10, mean ± SEM = 380.5000 ± 37.5287. Saline Day 2: n = 10, mean ± SEM = 447.5500 ± 34.8148. MSrAb + hS Day 0: n = 10, mean ± SEM = 376.0333 ± 32.6798. MSrAb + hS Day 2: n = 10, mean ± SEM = 360.6167 ± 42.7964. In **G**, Repeated measures ANOVA: Group (F (1, 10) = 0.0775, p = 0.7864), Time (F (1, 10) = 18.8533, p = 0.0015), Interaction (F (1, 10) = 0.1132, p = 7435). Vehicle Day 0: n = 6, mean ± SEM = 98 ± 13.4338. Vehicle Day 2: n = 6, mean ± SEM = 122.3333 ± 20.4999. BTKi Day 0: n = 6, mean ± SEM = 94.6667 ± 7.7230. BTKi Day 2: n = 6, mean ± SEM = 115.5 ± 7.7707. Post-hoc t test with Bonferroni correction: Vehicle Day 0 vs. Day 2 (F (10) = 3.3082, p = 0.0158), BTKi Day 0 vs. Day 2 (F (10) = 2.8324, p = 0.0356). In **H**, Two-way ANOVA: Group (F (1, 18) = 0.0121, p = 0.9135), Time (F (1, 18) = 34.9800, p < 0.0001), Interaction (F (1, 18) = 0.0270, p = 0.8712). Vehicle Day 0: n = 7, mean ± SEM = 651.2152 ± 68.3296. Vehicle Day 2: n = 7, mean ± SEM = 2181.1625 ± 248.9160. BTKi Day 0: n = 4, mean ± SEM = 720.3179 ± 145.7748. BTKi Day 2: n = 4, mean ± SEM = 2167.4881 ± 478.8819. Post-hoc Bonferroni's: Vehicle Day 0 vs. Day 2 (F (18) = 5.04, p = 0.0002), BTKi Day 0 vs. Day 2 (F (18) = 3.604, p = 0.0041). In **I**, Two-way ANOVA: Group (F (1, 17) = 0.0001, p = 0.9920), Time (F (1, 17) = 2.8110, p = 0.1119), Interaction (F (1, 17) = 1.0660, p = 0.3164). Vehicle Day 0: n = 6, mean ± SEM = 380.1389 ± 52.5780. Vehicle Day 2: n = 7, mean ± SEM = 341.3810 ± 44.5797. BTKi Day 0: n = 4, mean ± SEM = 441.6250 ± 89.5967. BTKi Day 2: n = 4, mean ± SEM = 278.6667 ± 58.6289. * p < 0.05, ** p < 0.01, *** p < 0.001, n = mice (large dots); up to 3 cells (small dots) were averaged per mouse in C, F, H, I; two-sided statistical tests. See **Extended Data Table 1** for statistical details.

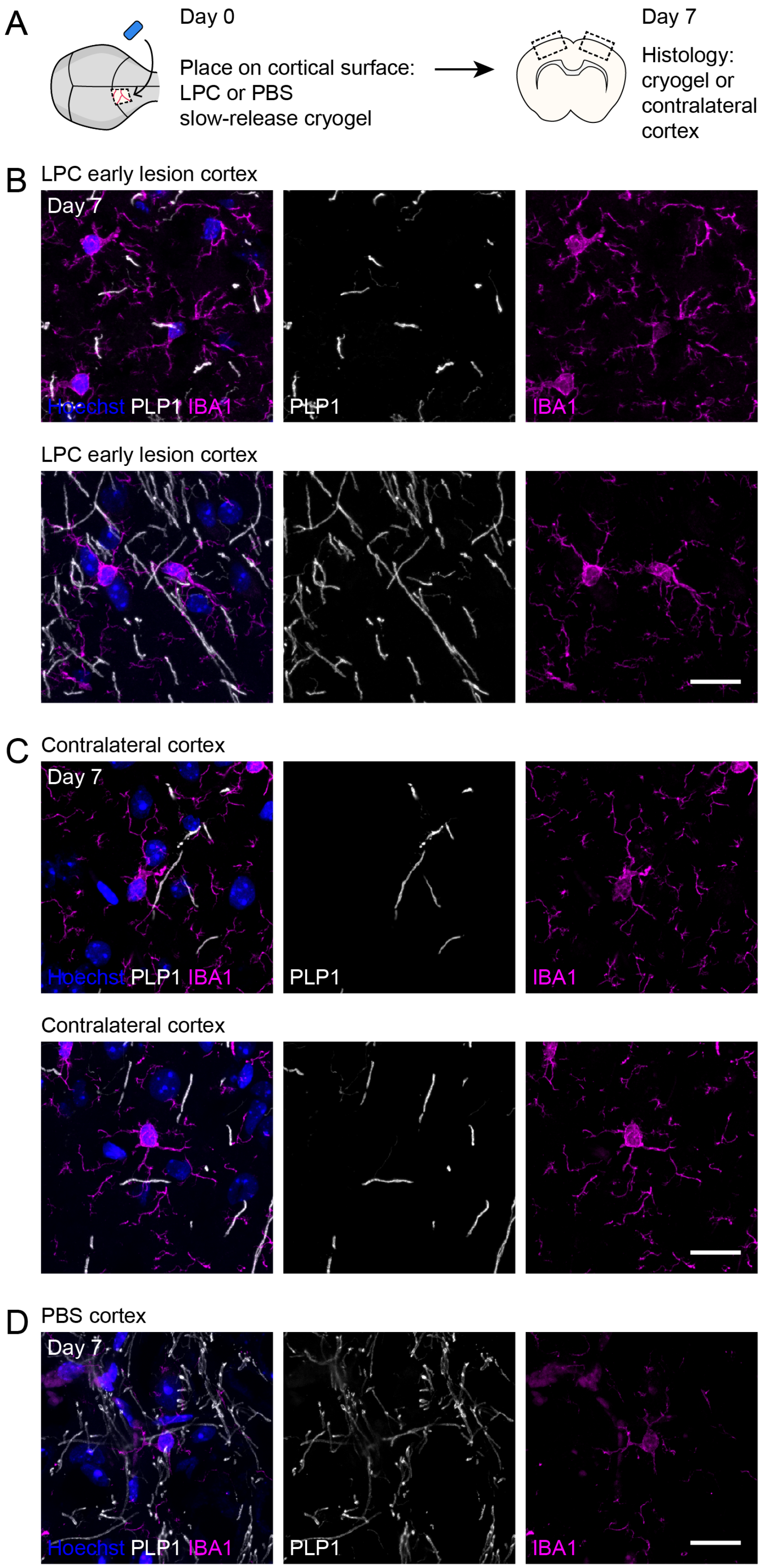
A
Day 0
Place on cortical surface:
LPC or PBS
slow-release cryogel
Day 7
Histology:
cryogel or
contralateral
cortex
B
LPC early lesion cortex
Day 7
Hoechst PLP1 IBA1
PLP1
IBA1
LPC early lesion cortex
C
Contralateral cortex
Day 7
Hoechst PLP1 IBA1
PLP1
IBA1
Contralateral cortex
D
PBS cortex
Day 7
Hoechst PLP1 IBA1
PLP1
IBA1

**Extended Data Fig. 6: Toxin-mediated demyelination does not induce microglia envelopment. (A)** Experimental timeline for LPC or PBS cryogel application and histology. Histology was timed to correspond to early demyelination. **(B-D)** Representative images of myelin (white) and microglia (magenta) in LPC early lesion cortex (B), cortex contralateral to LPC cryogel (C), and cortex underlying the PBS cryogel (D). Lesion area in LPC cryogel cortex was determined by increased numbers of microglia at this early demyelination timepoint. No evidence of microglial envelopment of myelin sheaths was observed. Scale bars are 20 μm. LPC = L-α-Lysophosphatidylcholine; PBS = phosphate buffered saline.

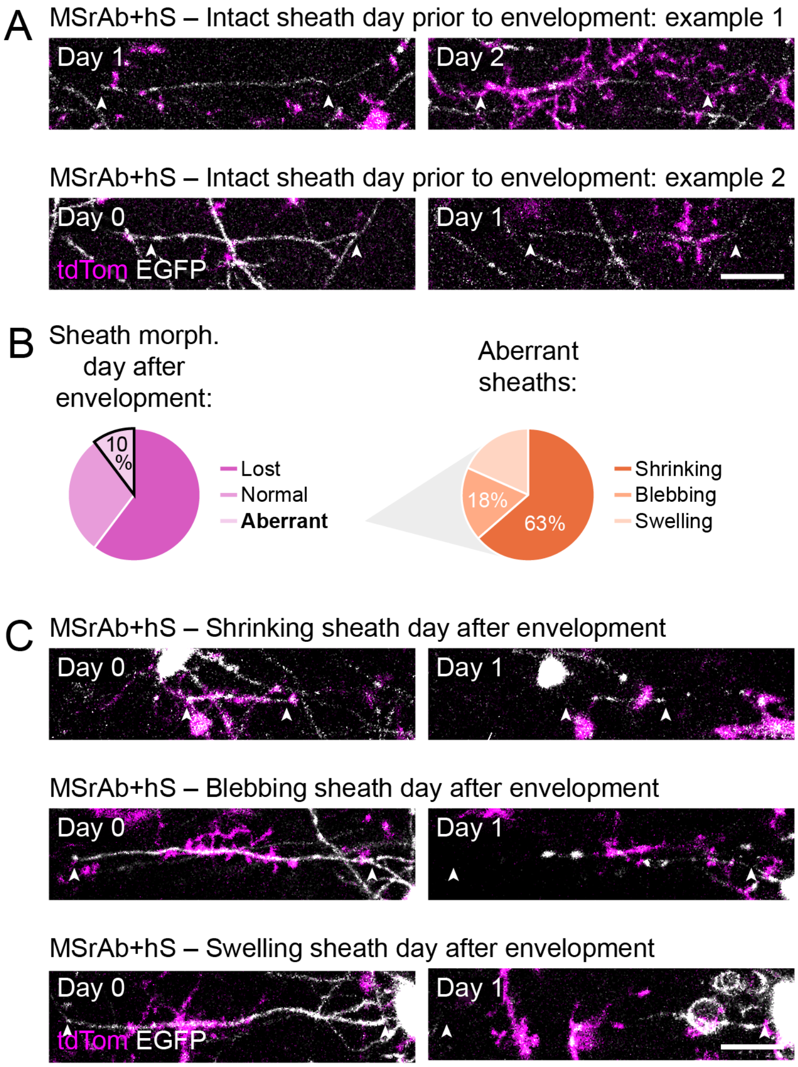


**Extended Data Fig. 7: Myelin sheath morphology before and after microglia envelopment. (A)** Example longitudinal *in vivo* images showing the typical normal morphology of sheaths (white) at the timepoint prior to their envelopment by microglia (magenta) in MSrAb + hS treated mice. Arrowheads point to both termini of sheaths. Scale bar is 20 μm. **(B)** Sheaths assuming an aberrant morphology the day after microglia envelopment were either shrinking (63%), blebbing (18%), or swelling (18%). N = 11 aberrant sheaths from 5 MSrAb + hS treated mice. **(C)** Example longitudinal *in vivo* images showing shrinking, blebbing, and swelling morphology of sheaths (white) at the timepoint after their envelopment by microglia (magenta) in MSrAb + hS treated mice. Arrowheads point to both termini of sheaths at first timepoint. Scale bar is 20 μm.

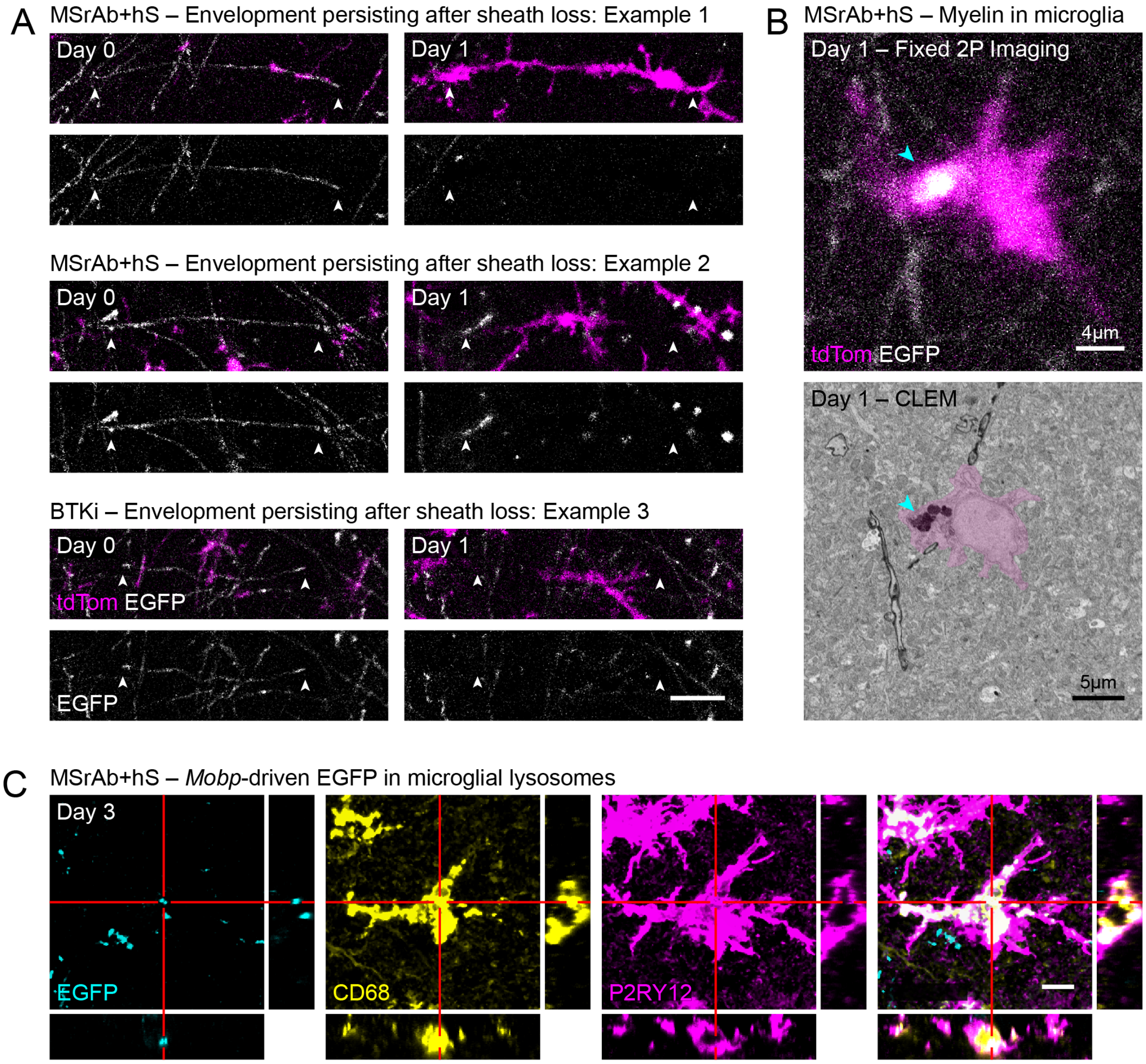


**Extended Data Fig. 8: Microglial phagocytosis of myelin in MSrAb demyelination model. (A)** Example longitudinal *in vivo* images of microglial envelopments (magenta) with a phagocytic-like morphology persisting after underlying myelin sheath (white) is lost in MSrAb + hS treated mice or those also given BTKi. For clarity, images were processed to remove bleed-through from the red fluorophore (microglia) into the green (myelin) channel. Arrowheads point to both termini of lost sheaths. Scale bar is 20 μm. **(B)** *In vivo* image of a microglia (magenta) with an EGFP (white) inclusion (arrowhead) on Day 1 (top). Electron micrograph of same microglia shows this inclusion (arrowhead) is electron-dense myelin-like debris (bottom). **(C)** Example image of immunofluorescence for EGFP+ myelin/oligodendrocytes (cyan), CD68+ lysosomes/endosomes (yellow), and P2RY12+ microglia (magenta) at Day 3 shows *Mobp*-driven EGFP inside microglial lysosomes/endosomes. Red lines highlight phagocytosed EGFP of interest. For each channel: Main image is XY maximum intensity projection of 57 1 μm-Z slices. Scale bar is 10 μm. Right image is ZY maximum intensity projection of 7 0.24 μm-X slices. Bottom image is XZ maximum intensity projection of 7 0.24 μm-Y slices.

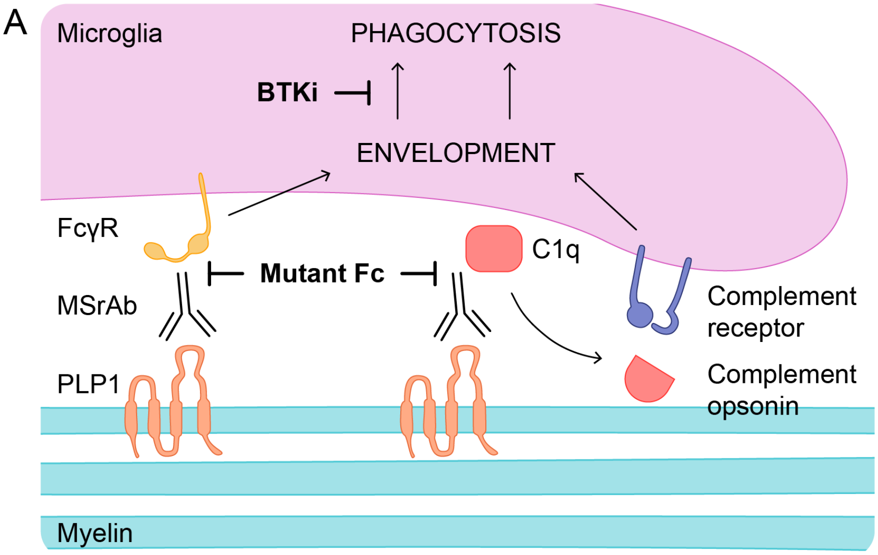


**Extended Data Fig. 9: Model of microglia-mediated demyelination. (A)** MS patient-derived antibodies target myelin protein PLP1 complexes[11]. Microglia recognize antibody via Fc gamma receptors or complement receptors, leading to sheath envelopment. Mutating the antibody Fc region to block Fc gamma receptor and complement C1q binding prevents microglia envelopment. Microglia then phagocytose enveloped sheaths, dependent on BTK signaling. Inhibiting BTK prevents the loss of sheaths following their envelopment.
